# Single-cell RNA sequencing identifies regulators of differentiation and nutritional cues in *Drosophila* female germ cells

**DOI:** 10.1101/2022.08.15.504025

**Authors:** Zhipeng Sun, Todd G. Nystul, Guohua Zhong

## Abstract

*Drosophila* ovarian germline stem cells (GSCs) are powerful model for stem cell research. However, due to the scarcity of GSCs in ovarian tissue, it is difficult to obtain the transcriptional profile of GSCs and identify novel GSC markers. In this study, we took advantage of single cell RNA sequencing (scRNA-seq) to profile the germline cells and somatic cells in wild type *Drosophila* ovary. We then performed an *in vivo* RNAi screen and network analysis to identify genes that are involved in the early stages of germ cell differentiation. We identified 33 genes with limited expression during early germ cell development and identified 19 genes that potentially regulate germ cell differentiation. Among these, an uncharacterized gene, which we named *eggplant* (*eggpl*), is highly expressed in GSC and downregulated in early daughter cells. Upon RNAi knockdown of *eggpl*, we observed an increase in germ cell proliferation, an accumulation of cysts in the early mitotic (2- and 4-cell) stages and an increase in overall ovary size compared to control when flies were maintained on a standard diet. In flies fed a rich yeast diet, the expression of *eggpl* was significantly lower and the effects of *eggpl* RNAi were suppressed, suggesting that downregulation of *eggpl* may link nutritional status to germ cell proliferation and differentiation. We also found that the matrix metalloproteases, *Mmp1* and *Mmp2* as well as the tissue inhibitor of metalloproteases (*Timp*) are additional regulators of *eggpl*. Collectively, this study provides new insight into the signaling networks that regulate early germ cell development and identifies *eggpl* as a key player in this process.

## 1. Introduction

The female reproductive system of *Drosophila melanogaster* has been well studied to understand the complex regulation of germline development (1, 2). Structurally, *Drosophila* ovary is made of 16-20 ovarioles, and 2-3 germline stem cells (GSC) reside in the anterior-most region of each ovariole (3). The GSCs reside in a specialized niche microenvironment and divide asymmetrically to produce one daughter cell which maintains the stem cell identity and another daughter cell that is displaced away from the niche and initiates differentiation as a cystoblast (CB). The CBs undergo four rounds of synchronous mitosis with incomplete cytokinesis, which eventually give rise to 15 nurse cells and 1 oocyte. Within the stem cell niche, several short-range extrinsic signals and intrinsic stemness-promoting factors are crucial to maintain the GSCs self-renewal and differentiation. The action of these signals is patterned by the somatic niche cells, which comprises terminal filament cells (TFs), cap cells (CpCs) and escort stem cells (ESCs) (4-6). *Decapentaplegic* (*dpp*) is one of the necessary and sufficient niche-derived factors for GSC maintenance. A high level of dpp signaling activity activates the bone morphogenic proteins (BMP) signaling pathway to transcriptionally silence the expression of differentiation promoting factor bag-of-marbles (bam), and sustains GSC identity. In contrast, CB positioned outside the niche experiences a weaker Dpp signal and triggers bam expression for differentiation (7). The escort cells (ECs) also express Epidermal Growth Factor (EGF) to promote a differentiation program in CB by repress the transcription of *division abnormally delayed* (*dally*) (8). In addition, both nuclear organization and chromatin modification are also play a key role in the regulation of GSC homeostasis. For example, it was reported that a linker histone H1 is intrinsically required for GSC maintenance, since the depletion of H1 in the germline cells would lead to premature expression of Bam and the loss of GSCs (9). Similar to *scrawny* (*scny*), an H2B ubiquitin protease, it is highly expressed in GSCs to suppress methylation at lysine residues and functionally repress target genes. Loss of *scny* results in early expression of Bam (10). It has also been shown that *dSETDB1* or *eggless* (*egg*), a histone methyltransferase, was responsible for GSC fate. Depletion of *egg* in GSC impairs self-renewal, while the *egg*-deficient GSCs could differentiate normally (11). In addition, stonewall (Stwl), a chromatin-associated protein which acts as a dominant Suppressor of variegation, is enriched in GSCs (12). Mei-P26 suppresses transcripts that promote differentiation in CBs by antagonizing miRNA pathway. However, *zpg* is essential to activate the differentiation of GSC progeny (13). Therefore, the proper balance of intrinsic and extrinsic gene expression is imperative for GSC self-renewal and differentiation.

An additional layer of regulation of the GSC niche comes from signals that communicate the availability of a rich diet. On high protein diet, GSCs and their descendent exhibit an increased rate of division and differentiation, and this response to diet is regulated by the evolutionarily conserved insulin signaling pathway. Neurosecretory cells in the brain produce the insulin-like peptides (DILPs), which directly regulate the G2 phase of GSC division and stimulate cyst growth (14-15). In flies fed a yeast-rich diet, the ovary size and egg production are significantly increased. (16). This is due to the action of insulin on the GSC niche cells, which facilitates GSC proliferation and maintenance, in part by promoting the extension of escort cell membranes to wrap around GSC and cysts (17). The membrane extensions are regulated by a membrane protein, Failed axon connections (Fax), which is induced by S6K activation downstream of the insulin receptor. Insulin also acts on cap cells to promote Notch Signaling and stimulate the physical adhesion between cap cells and GSCs through E-cadherin (18). However, the downstream response in germ cells to these nutrient-activated cues is not well understood.

In this study, we identified 33 genes that are differentially expressed during early germ cell development and that 19 of these are required for germ cell function. These genes were identified by scRNA-seq analysis of adult wildtype ovaries followed by validation of expression patterns *in vivo* and an RNAi screen for GSC decrease/increase, GSC loss and tumor formation. In addition, network analysis of the differentially expressed genes in undifferentiated germ cell-1 and -2 clusters revealed several common nodes. Among the genes we identified *CG32814*, is an uncharacterized gene which we have renamed *eggplant* (*eggpl*). We found that *eggpl* is specifically expressed in GSC at transcriptional level, but Eggpl protein is detectable in germ cells throughout Region 1. We find that *eggpl* is also expressed in larval male and female gonads and in adult testes as well. We also found that cell cycle in germ cell cysts was accelerated and the ovaries were larger in flies in which *eggpl* was depleted, either by RNAi or in a CRISPR knockout. In contrast, in flies fed a rich yeast diet, *eggpl* expression was reduced and the difference in germ cell proliferation rates and ovary size between the control and *eggpl* knockdown genotypes was reduced. Notably, we found that the MMP-dependent Timp pathway is an additional regulator of *eggpl* in germ cells. Taken together, these findings reveal the regulators that controls early germ cell differentiation and coordinate the rate of germline stem cell division with nutrient availability.

## 2. Materials and Methods

### 2.1. Fly stocks and fly husbandry

The gene names, genetic symbols, and detailed information about fly strains applied in this study are presented in the text and in FlyBase. All fly stocks were maintained at 25°C and reared on standard cornmeal agar food. For RNAi experiments, crosses were set up at 18°C and adults were transferred to 29°C upon eclosion for 7-9 days. The following flies were used in this study: *y*^*1*^*w*^*1118*^, *nos-Gal4/CyO;tub-Gal80*^*ts*^*/TB, vasa-EGFP, nos-Gal4, UAS*-*Timp, UAS*-*Timp*-*RNAi, UAS*-*Mmp1*-*RNAi, UAS-Mmp1·f2*^*E225A*^, *UAS-Mmp1·fDN*^*P10.PX*^, and *UAS-Mmp2-AGPI* (gifts from Suning Liu, South China Normal University, China), *bam-GFP* (a gift from Yu. Cai, Temasek Life Sciences Laboratory, Singapore).

The RNAi fly lines were obtain from Bloomington Stock center or Tsinghua Fly center and a full list of genotypes is provided as Supplementary Information (**Supplementary Table 2**).

### 2.2. Ovarian cell suspension for scRNA-seq

Newly emerged virgin female flies were fed for 1 week to encourage ovarian growth, then 150 flies were dissected in a petri dish containing 1mL of S-FBS (Serum-free Schneider’s insect medium (Sigma-Aldrich, cat. no. S0146) supplemented with 10% (v/v) fetal bovine serum (FBS), heat inactivated (Sigma-Aldrich, cat. no. F4135)) under the microscope (Leica, SAPO, Germany). After dissection, we discarded S-FBS and added 1 mL of PBS to rinse ovaries in a 1.5-ml centrifuge tube, and allowed samples to settle for 5 min, and then rinsed them twice with PBS. Dissociation was carried out at room temperature in 700 μl of dissociation medium by adding 70 μl of 5% (w/v) trypsin (Invitrogen, cat. no. 27250-018) and 70 μl of 2.5% (w/v) collagenase (Invitrogen, cat. no. 17018029) to 560 μl of PBS, and incubated for 15 min with continuous shaking. After incubation, the ovarian cell suspension was pipette into a 40-μm mesh cell strainer, and filtered suspension into a 1.5-ml centrifuge tube containing 500 μl of S-FBS. Then, the empty tube and cell strainer were washed by 100 μl of S-FBS respectively to collect the remaining cells. The cell suspension was collected by centrifuging 5 min at 425 × g, 4°C, and discarded the S-FBS and resuspended the pellets in each tube with 200 μl of serum-free Schneider’s insect medium, after that we combined the suspensions into one tube. Following, the cell viability was examined by using 0.4% trypan blue (Solarbio, cat. no. T8070) in the proportion of 1:1, and counted by a hemocytometer. The concentration of cell suspension was 1.37×10^6^ cells/ ml, and the viability was 90% at least, according to 10× Genomics recommendations.

### 2.3. Single-cell RNA sequencing

Single-cell libraries were constructed using Chromium single-cell 3’ Library (v2) kit via End Repair, A-tailing, Adaptor Ligation, and PCR according to the manufacturer’s protocol. In brief, the cells of each group were mixed into one sample and adjusted to 1000 cell/μl. Then, the indexed sequencing libraries which contained the P5 and P7 primers were prepared using Chromium single-cell 3’ Reagent kit, and the barcoded sequencing libraries were quantified using a standard curve-based qPCR assay (KAPA Biosystems, USA) and Agilent Bioanalyzer 2100 (Agilent, Loveland, CO, USA). Subsequently, the library sequencing was performed by Illumina HiSeq 4000 with a custom paired-end sequencing mode 26 bp (read 1) × 98 bp (read 2).

### 2.4. 10×Genomics initial quality control

The scRNA-seq data were processed with the Cell Ranger Single Cell Software Suite (v6.1) (http://software.10xgenomics.com/single-cell/overview/welcome) for quality control, sample demultiplexing, barcode processing, and single-cell 3’ gene counting. First, the raw data were demultiplexed by using an 8 bp index read at the end of Read 1 and Read 2 paired-end reads, to generate FASTQ files, and then quality control was performed using FastQC, and these data were aligned against the Nucleotide Sequence Database (https://www.ncbi.nlm.nih.gov/genbank/) using the NCBI Basic Local Alignment Search Tool (BLAST). Second, the reads were aligned to the *Drosophila* reference genome (dm6) (https://www.ncbi.nlm.nih.gov/assembly/GCF_000001215.4#/st) by STAR RNA-Seq aligner. Once aligned, barcodes associated with these reads-UMIs were subjected to filtering and correction. For UMI tag counting, the 10× Genomics pipeline Cell Ranger was used to generate single-cell gene counts for each library. The confidently mapped, non-PCR duplicates with valid barcodes and UMIs were eventually used to generate the gene-barcode matrix. For the higher-depth libraries, the samples were normalized to the sample sequencing depth. CellRanger version 2.0.0 and Seurat (v4.0.4) (19). R package were used to filter out the low-quality cells, and the following criteria were used to filter cells: (1) gene counts >3000 per cell; (2) UMI counts >12 000 per cell; and (3) percentage of mitochondrial genes >30%. In thisstudy, the estimated cell number was derived by plotting the UMI counts against the barcodes and revealed 21755 cells used for downstream analysis. Based on the transcriptomes of 21755 cells, a total of 0.39 billion clean reads achieving an average read of 18202 per cell and the ratio of high-quality reads to qualify scores at Q30 was more than 90.6% were obtained. The total number of read pairs that were assigned to this library in demultiplexing is 395,988,785, and the valid Barcodes (Fraction of reads with barcodes that match the whitelist after barcode correction) and valid UMIs (unique molecular identiifier) are 97.8% and 100% respectively. The number of estimated cells is 21,755 with 18,202 mean reads per cell, and the number of median genes per cell was 638. The rough sequencing data were filtered according to the criterion that any cell containing more than 25,000 UMIs counts and more than 30% mitochondrial UMIs was filtered out. We finally obtained 8497 out of 21,755 cells with 3,993 median UMIs per cell and 868 median genes per cell (**Supplementary Table. 1**) for scRNA-seq analysis.

### 2.5. Clustering analysis

For the clustering, we used principal component analysis (PCA) to normalize and filter the gene-barcode matrix and to reduce feature dimensions. The top 5 major components were selected to obtain the visualized 2D clustering image using T-distributed stochastic neighbor embedding (tSNE). The graph-based clustering method was applied to group cells with similar expression patterns of marker genes. The ovarian cell clusters were grouped into 24 unsupervised categories using the different resolution parameters (R=0.5 or default values). The pairwise Pearson correlation was calculated between each cluster for hierarchical clustering. Based on the differentially expressed gene results, a visualized heat map was created using Seurat (v4.0.4) R package. The tSNE plot was generated for a graphical representation of specific gene expression by Loupe Cell Browser software and Seurat (v4.0.4) R package. Notably, in order to improve the accuracy of trajectory based on our clustering results, we removed the cells which expressed somatic cell marker *tj*, and non-*vasa* (*vas*) expressing cells in germline clusters.

### 2.6. Marker gene analysis and Monocle pseudotime analysis

The candidate marker genes which enriched in a specific cluster were selected according to the expression profile of top genes among 24 clusters, and the putative biological identity of each cluster was assigned based on the expression patterns of highly expressed genes and experimentally validated markers. Single-cell pseudotime analysis were carried out by using matrices of cells and gene expression by Monocle (v2.20.0) which provided the visualized trajectory with tips and branches in the reduced dimensional space.

### 2.7. Differential gene expression analysis

The likelihood-ratio test (20) was used to seek differential expression profiles in each cluster, and the following criterion were allowed to identify the differentially expressed genes: (1) P-value ≤ 0.01. (2) Log2(fold change [FC]) ≥ 0.360674. (3) The percentage of cells where the gene is detected in a specific cluster >25%. Then, Gene Ontology (GO) enrichment analysis was performed to filter the differentially expressed genes that correspond to biological functions. The peak-related genes were mapped to GO terms in the GO database (http://www.geneontology.org/), and the significantly enriched GO terms were defined by a hypergeometric test. To further understand the biological functions of these genes, we used Kyoto Encyclopedia of Genes and Genomes (KEGG) (https://www.kegg.jp/) pathway enrichment analysis to identify the enriched metabolic pathways and signal transduction pathways.

### 2.8. Gene regulatory network analysis

The transcription factor network inference was conducted by SCENIC R package. The log-normalized expression was generated by using Seurat, and the pipeline was implanted step by step. Preliminarily, the gene co-expression was identified via GENIE3, which may include some false positives and indirect targets. Then, we identified putative direct-binding targets by pruning each co-expression module via Rcis Target. Precisely, networks (regulons) were retained if the TF-binding motif was enriched among its targets, while target genes without direct TF-binding motifs were removed. Last, we scored the activity of each regulon for each single cell via the AUC scores using AUCell R package.

The *Cytoscape* v.3.9.1 software was applied for the construction of gene regulatory network according to its online user manual.

### 2.9. RNA in situ hybridization

Probe synthesis. Using the genomic DNA as a template, and amplifying the exon regions of targeted genes by using the primers with SP6 sequence (ATTTAGGTGACACTATAGAAGNG) according to the product description of KAPA HiFi PCR Kit (Roche Diagnostics, cat. no. 07958927001). Sense and antisense digoxigenin (DIG)-labeled probes were synthesized from the purified PCR product using DIG RNA Labeling Kit (Roche, cat. no.11175025910). All primer sequences were listed in Supplementary Information (Supplementary Table 3).

The procedures for RNA *in situ* hybridization were as follows. Briefly, the samples were dissected in PBS and immediately fixed in 4% PFA with 0.1 M Hepes at 4°C overnight. On the next day, the samples were washed 3 × 10 min with PBST (0.1% Tween 20 in PBS) and dehydrated with sequential washes with 50% and 100% methanol in PBST for 5 min each time. Then, the samples were stored in the -20°C refrigerator for 40 min, and washed with PBST 3 × 10 min before proteinase K (Sigma-Aldrich, cat. no. 39450016) treatment for 5 min at room temperature. Samples were washed with PBS for 5 min and fixed with 4% PFA for 20 min, then washed with PBST 3 × 10 min and incubated in hybridization buffer (50% formamide, 5x SSC, 0.1% Tween-20, 50 µg/µl heparin, and 100 µg/ml salmon sperm DNA) with probe in hybridization oven (Jingxin industrial development co. ltd, LF-I) at 60°C for 24 h at least. After hybridization, the samples were washed 4 × 30 min at 60°C, once with 2x SSCT (2x SSC, 0.1% Tween-20) for 15 min and twice with 0.2x SSCT (0.2x SSC, 0.1% Tween-20) for 30 min each at 60°C. Next, samples were washed with MABT (0.1 M maleic acid; 0.15M NaCl PH 7.4 and 0.1% Tween-20) 2 × 10 min at room temperature and blocked for at least 30 min, and then added anti-dig-POD (1:200; Roche, cat. no.11207733910) in 5% blocking solution (Roche, cat. no.11096176001) at room temperature overnight. Finally, the fluorescence reaction was carried out by using TSA fluorescein system (Perkin Elmer, cat. no. TS-000100) for 1.5 h in dark and subsequently used Hoechst 33258 (Sigma-Aldrich, cat. no. 23491454) to label the nucleus.

### 2.10. Construction of the transgenic fly lines

We first designed guide RNA targets with: 1. Chopchop (https://chopchop.cbu.uib.no/) (21), 2. CCTop (https://cctop.cos.uni-heidelberg.de/) (22). Genomic DNA was isolated from the injection stock. PCR was performed using primers flanking the targets. The amplified products were sent for Sanger sequencing. If SNPs were found on the targets, gRNA sequence would be modified to be consistent with the target sequence of the stock.

The first base of gRNA sequence was changed to G for the T7 transcription. Following the protocol (23), template for in vitro transcription by T7 polymerase was generated by annealing of two DNA oligonucleotides and PCR amplification. *In vitro* transcription was performed with the T7 RiboMAX™ Kit (Promega, cat. no. P1320). Transcripts were purified by phenol-choloroform extraction and isopropanol precipitation.

Plasmid MLM3613 (Addgene plasmid, cat. no. 42251) was linearized with Pme I (New England Biolabs) and purified by ethanol precipitation. Cas9 mRNA was transcribed with mMESSAGE mMACHINE® T7 Transcription Kit (Ambion, cat. no. C013843), polyadenylated with the E.coli Poly (A) polymerase Kit (NEB, cat. no. M0276L), and purified with the RNeasy Mini Kit (QIAGEN, cat. no. 74106).

To knock in the 6 × HA of GFP in the N-terminal of *eggpl*, the pBluescirpt SK vector (pBS) was used as the backbone. Using genomic DNA of the injection stock, the homology 5’arm and 3’arm was amplified and linked to the pBS backbone with Gibson Assembly Kit (NEB, cat. no.E2611L) as ‘pBS-CG32814-arm’. Then the pBS-eggpl-arm was linearized by PCR and linked to the GFP-6HA cassette with Gibson Assembly Kit (Thermo Fisher, cat. no. A46624), and that produced the final donor construct ‘pBS-CG32814-GFP-6HA’ (supplementary Fig.3 A).

To generate a mutant allele of *eggpl*, we used Cas9/CRISPR to introduce mutations downstream of the ATG in the *eggpl* open reading frame. We identified an allele, *eggpl*^*[1]*^ in which the 5 base pairs immediately downstream from the ATG (`AGTAG`) were deleted and a 28-base pair region that is 50 base pairs downstream from the ATG (TTAAAACGGACACCATCGGCGAAGAAAA) contained multiple deletions and substitutions. In *eggpl*^*[1]*^, this 28 base pair region was instead an 18 base pair region with the following sequence: CTTCTTCACCATTTTCAC. The resulting allele contains several frameshifts that alter the coding sequence of the 5’ end of the gene but restore the normal reading frame after the 27^th^ codon. The DNA sequence of this mutated region in the *eggpl*^*[1]*^ allele, starting at the ATG of the open reading frame, is ATG-----CGGAATCATTTTCAGACAGAATCCAGATGGATCTTTTTCACCTTCTt---C-ttCACCATtttC------AcC. Dashes indicate the location of deletions and lowercase letters indicate substitutions, relative to the wildtype sequence. The gRNA sequences is list below,

CG32814-sg1: GTTAGAATCAAAATGAGTAG

CG32814-sg2: GCTTTTAAAACGGACACCAT

The primers for validation was GGAGTCTCCAGCAATTACTGTAT and CCCTGATTGCAATGAGTTTTCAGT.

15ug of Cas9 mRNA and 7.5ug sgRNA were mixed with DEPC water in a 30ul volume. And the RNA mix injection was performed by Qidong Fungene Biotechnology (http://www.fungene.tech). 300 embryos were injected.

When the injected P0 embryos grew into adults, they were crossed with Fm7a. The genomic DNA of the P0 and F1 flies was extracted. PCR was performed using primers for validation of mRuby3 insertion:

F: AAGTTGTCAGCCGATTGGCGTGG

R: ATTCACTTTTCCATTATTGAATG

The F2 flies from positive F1 tubes were balanced with *Fm7a*.

Transgenic fly lines w*; P{UAS-eggpl-GFP}attP2/TM6B was generated by integrating *UAS-eggpl-GFP* into the attP2 site. Briefly, the NotI/XbaI PCR fragment of eggpl-CDS (GFP tag) was cloned into the NotI/XbaI sites of pJFRC28-10 × UAS-IVS-GFP-p10 vector (Addgene Plasmid, cat. no. 36431). The primer pairs used for PCR validation were as following:

eggpl-3F: ACAACAAAGCCATATGATGAG

p10-R: GCCACTAGCTCGCTATACACT

### 2.11. Whole-mount immunofluorescence staining and confocal imaging

Ovaries were dissected in PBS and fixed in 4% PFA, 0.1 M Hepes, PH 7.4 for 30 min at room temperature with gentle rotation, then washed the ovaries 3 × 15 min with 500 μL 0.1% PBT (0.1% Triton X-100 in PBS). The samples were blocked in 5% NGS buffer (5% normal goat serum in 0.1% PBT) for 1 h before incubation with primary antibody at room temperature overnight. The following day, diluted primary antibody was collected for reuse, and the samples were washed 3 × 15 min with 500 μL PBT and incubated with diluted secondary antibody for 3h with rotation. The Hoechst labeling was performed after washing with PBT 3 × 15 min. Finally, the samples were mounted on slides in Vectashield mounting medium and stored at 4°C. Note that, 50% normal goat serum in 0.1% PBT and higher concentration of diluted primary antibody were recommended to apply for the continuous antibody staining procedure after RNA *in situ* hybridization. For pMad staining, the samples were suggested to fix in 4% PFA for 50 min, and washed the ovaries with 0.1% PBT three times for 3 h at least.

The following primary antibodies were applied in this study: mouse anti-α-Spectrin (3A9, 1:100; Developmental Studies Hybridoma Band (DSHB)), rabbit anti-pMad (1:800; Cell Signaling), chicken anti-GFP (1:5,000; Abcam), rat anti-Vasa (1:100, DSHB). Alexa Fluor 488, 555, or 633 conjugated goat secondary antibodies (gifts from Yu. Cai, Temasek Life Sciences Laboratory, Singapore) against mouse (1:500), rabbit (1:1,000), chicken (1:500) and rat (1:1,000) were used to detect the primary antibodies. Polyclonal anti-TfIIA-S and anti-eggpl were generated via immunization of rabbits (Custom made for chickens by GeneCreate Biotech Co., Wuhan, China, 1:1,000). The DNA dyes Hoechst 33258 (1:5,000; Cell Signaling Technology) was used to label the nucleus.

Images were captured by using the Nikon A1 plus confocal microscope (Nikon, Japan) with APO 60×/1.40 oil objective lens at room temperature, and all images were processed with NIS-Elements software for image acquisition and analysis. The mean of fluorescence intensity was examined by using ImageJ v1.8.0 software according to the manual instruction (supplementary Fig.3 C) or Imaris (Fig. 5L-M). For image analysis in Imaris, surfaces were generated in the green (GFP) channel and manually split or merged to generate a single surface for each cyst in Region 1. Then, the mean pixel intensity in the green channel within each surface was calculated.

**Fig. 1.**
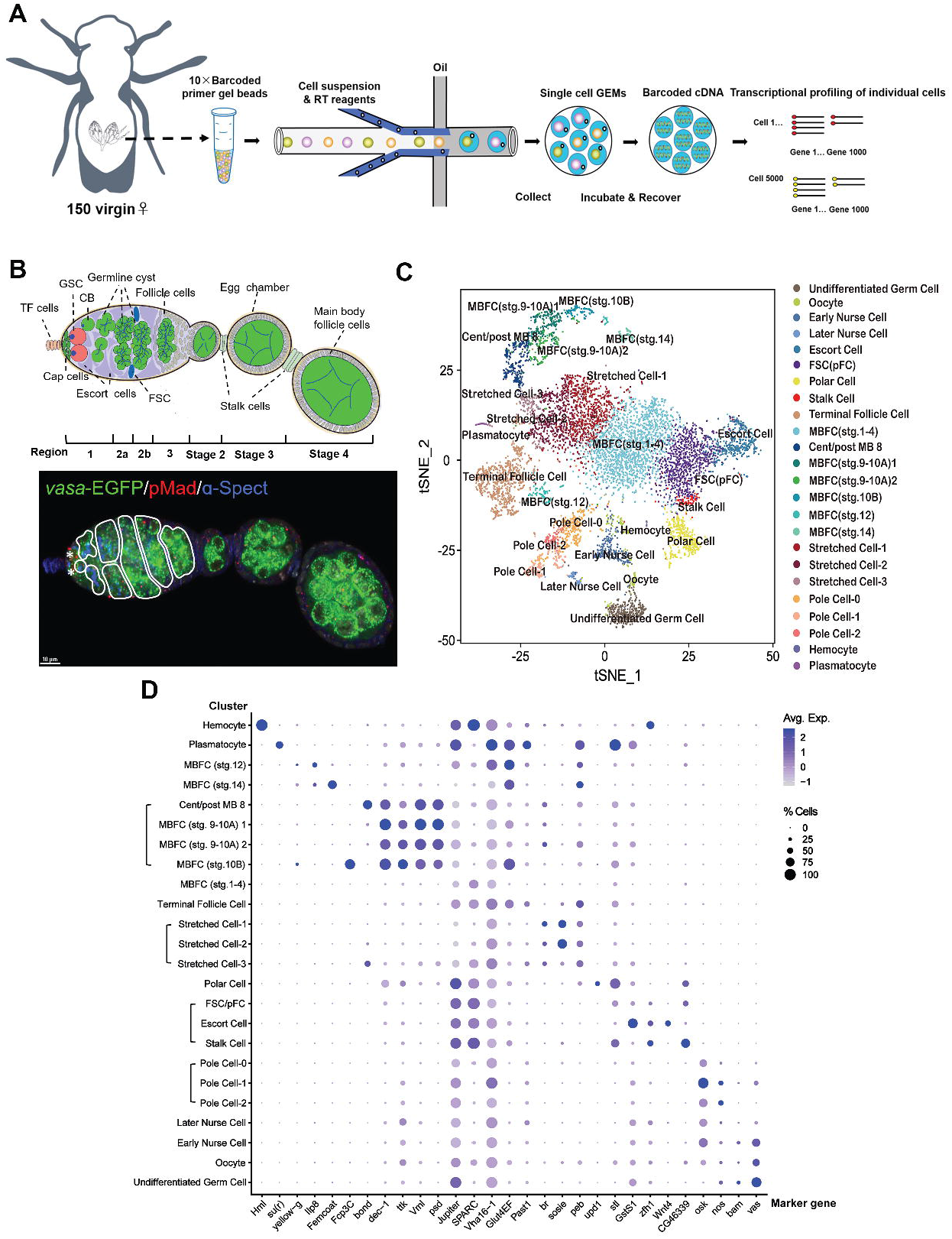
10×Genomics single-cell RNA sequencing on adult *Drosophila* ovary. **(A)** Schematic of experimental workflow for cell capture and single cell data analysis. **(B)** Illustration of a *Drosophila* ovariole, describing asymmetric divisions of GSCs and CB, which divide four times to produce developing germline cysts. By region 2b, 16-cell cyst was completely formed and surrounded by follicle cells. As the cyst moves down to region 3, the egg chamber containing 1 oocyte and 15 nurse cells is formed and ready to bud off. The *vasa-EGFP* line was used to visualize all germ cells along germline. Anti-pMad (red) was used to label GSCs (asterisk), while anti-α-Spectrin (blue) was used to stain spectrosomes (round dot) and branched fusomes. **(C)** The t-SNE plot of 24 distinct cell clusters marked with different colors. **(D)** The dot plot of scaled expression of selected typical marker genes in each cell type.

**Fig. 2.**
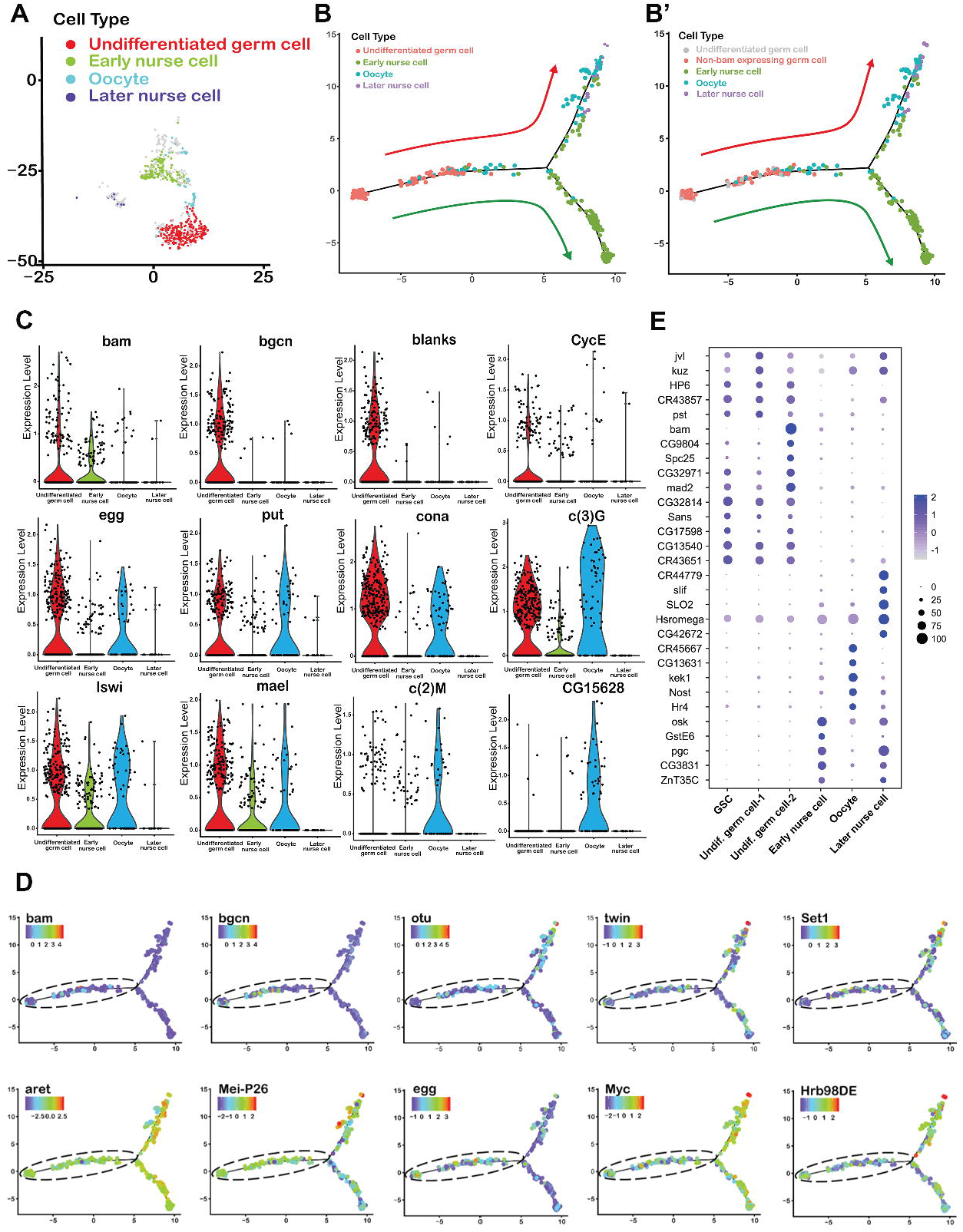
The identification of GSCs and germ cell subclusters. **(A)** tSNE plot revealing 4 germ cell subclusters, undifferentiated germ cells (red), early nurse cells (green), oocytes (blue) and later nurse cells (purple). **(B-B’)** The monocle analysis reveals the developmental linear trajectory of germ cells, and the putative GSCs population was distinguished in non-bam expressing germ cells which located in the beginning of trajectory. Arrows indicate the direction of differentiation. **(C)** To assign identities to these germ cell subclusters, the violin plots were used to visualize the distribution of normalized typical marker genes expression levels. **(D)** The expression of GSC maintenance genes (*aret, Mei-P26, egg, Myc* and *Hrb98DE*) and differentiation genes (*bam, bgcn, out, twin* and *Set1*) along the primary branch (dotted circle line) in pseudotime. **(E)** The dot plot presents the respective top 5 genes in the GSCs, undifferentiated germ cell-1, undifferentiated germ cell-2, early nurse cells, oocytes and later nurse cells. The dot diameter represents the percentage of cell expressing top genes.

**Fig. 3.**
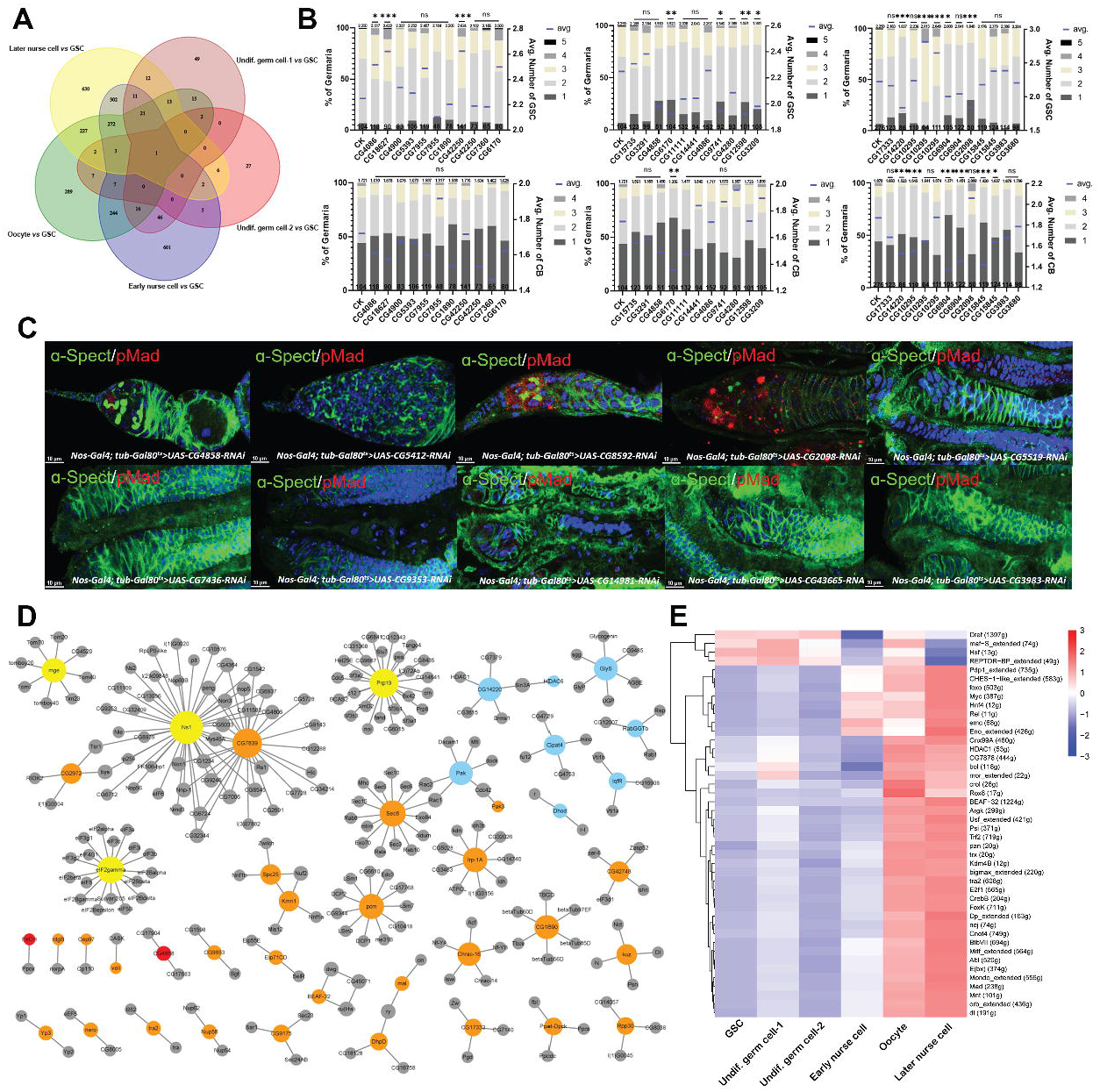
The construction of gene regulatory network in GSCs. **(A)** The Venn diagram of the number of differentially expressed gene. **(B)** The statistical analysis of the average number of GSC and CB in RNAi lines of screened differentially expressed gene. Error bars show SEM, ns indicates no significant difference, \**P*<0.05, \*\**P*<0.01, \*\*\**P*<0.001. **(C)** The typical phenotypes of germaria in selected RNAi lines stained with anti-α-Spectrin (green) and anti-pMad (red). **(D)** The construction of gene interaction network by using *Cytoscape 3.9.1* software. **(E)** The heat map of SCENIC analysis on GSC, undifferentiated germ cell-1, undifferentiated germ cell-2, early nurse cell, oocyte and later nurse cell subclusters.

**Fig. 4.**
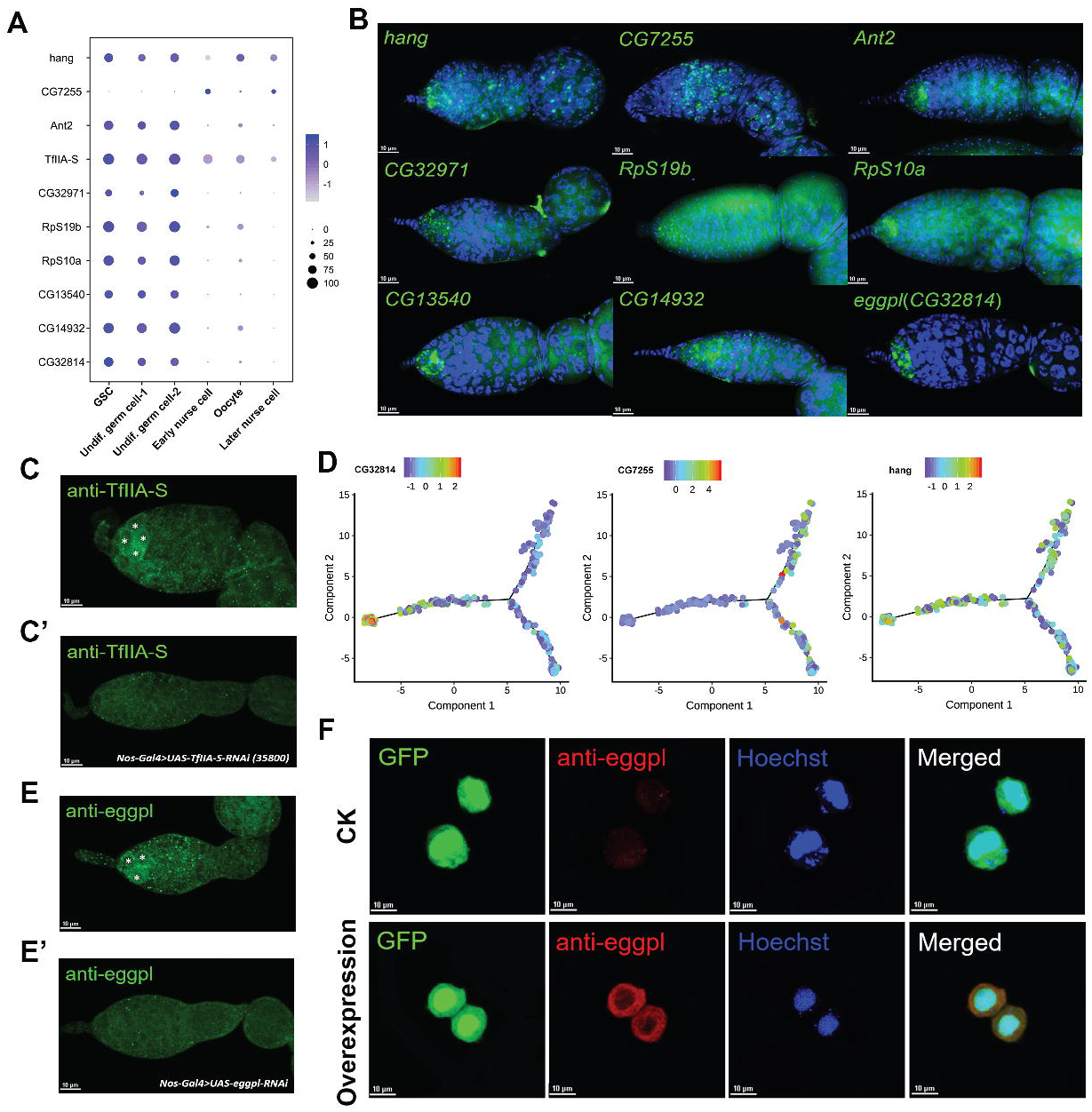
The validation of GSC marker genes. **(A)** The dot plot showing the expression of selected specific marker genes in each subclusters. The color intensity from dark to light represents the average normalized gene expression level. **(B)** The expression patterns of candidate marker genes was validated by using *in situ hybridization*. **(C-C’)** Immunofluorescence staining with anti-TfIIA-S on wild type and *nos-Gal4>UAS-TfIIA-S-RNAi* (negative control). **(D)** The expression profiles of *eggpl, CG7255* and *hang* along the trajectory branches in pseudotime. **(E-E’)** Immunofluorescence staining with anti-eggpl on wild type and *nos-Gal4>UAS-eggpl-RNAi* (negative control) ovary. **(F)** The overexpression of GFP (green) and eggpl in *Sf9* cell line *in vitro*. The anti-eggpl (red) was used to detect the eggpl protein, and Hoechst (blue) was used to label the nucleus.

**Fig. 5.**
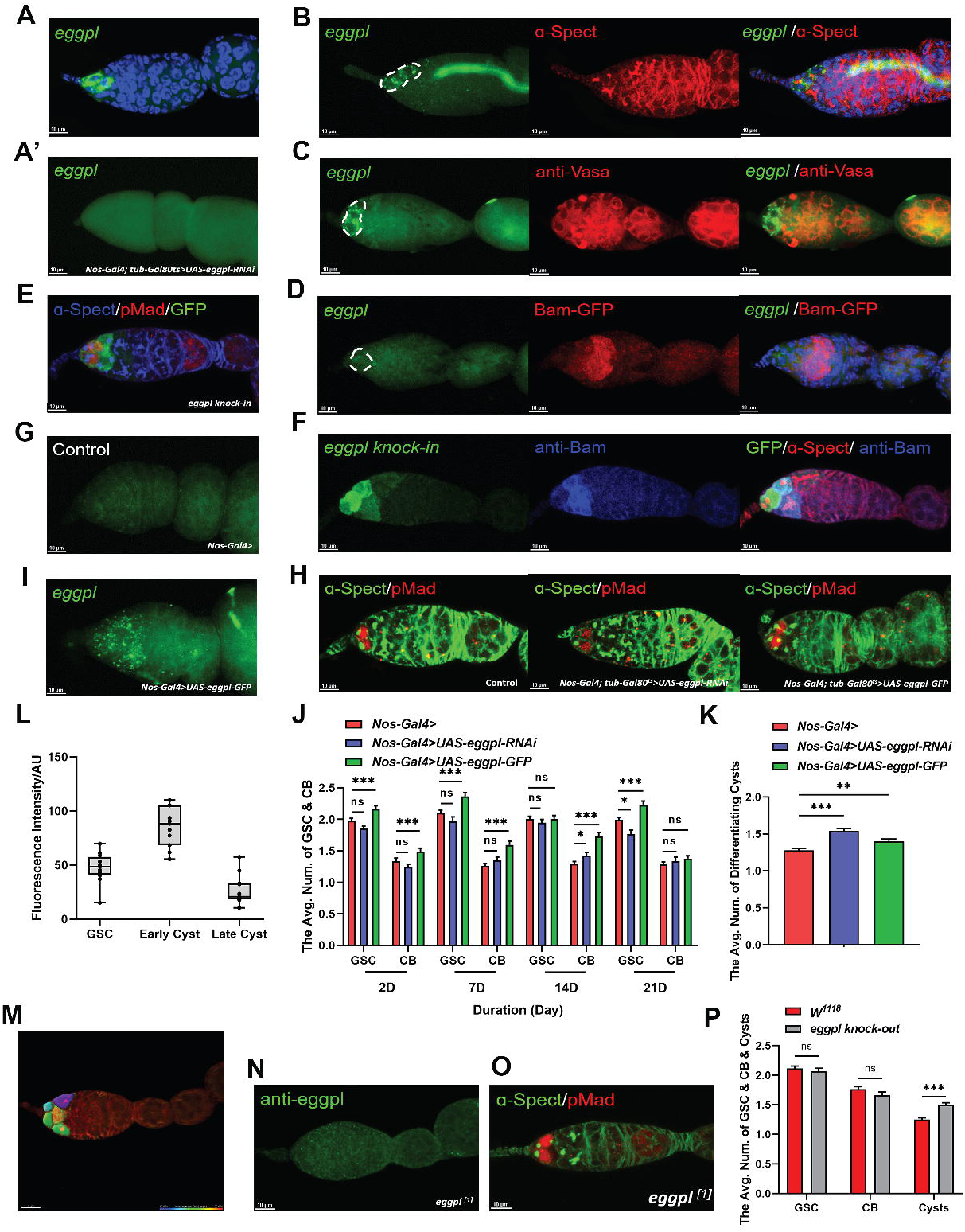
The characteristics of *eggpl* expression in ovarian germline. **(A-A’)** The mRNA *in situ hybridization* of *eggpl* (green) on wild type and *nos-Gal4>UAS-eggpl-RNAi* (negative control) line. **(B-C)** Immunofluorescence staining with anti-α-Spectrin (red) and anti-Vasa (red) on *eggpl*-*in situ* (green) labelled tissues respectively. **(D)** The mRNA *in situ hybridization* of *eggpl* (green) on *bam-GFP* line stained with GFP (red). **(E)** Immunofluorescence staining with anti-α-Spectrin (blue), anti-pMad (red) and GFP (green) on the *eggpl::GFP knock-in* line. **(F)** The anti-GFP, anti-α-Spectrin (red) and anti-Bam (blue) were used to stain on *eggpl::GFP knock-in* line. **(G and I)** The mRNA *in situ hybridization* of *eggpl* (green) on *nos-Gal4>* (negative control) and *nos-Gal4>UAS-eggpl-GFP* lines. **(H)** The phenotypes of wild type, *nos-Gal4>UAS-eggpl-RNAi* line and *nos-Gal4>UAS-eggpl::GFP* line staining with anti-α-Spectrin (green), anti-pMad (red). **(J)** The average number of GSC and CB in *nos-Gal4* line, *nos-Gal4>UAS-eggpl-RNAi* line and *nos-Gal4>UAS-eggpl::GFP* line on 2 day, 7 day, 14 day and 21 day. **(K)** The average number of differentiating germline cysts in three types of fly lines. **(L)** Quantification of the GFP intensity mean in *eggpl::GFP knock-in* line, n=11. **(M)** Image illustrating the image segmentation used to quantify the expression pattern of *eggpl*. **(N)** Immunofluorescence staining of an *eggpl*^*[1]*^ germarium with anti-eggpl antibody showing a lack of signal. **(O)** The typical phenotype of *eggpl*^*[1]*^ germarium staining with anti-α-Spectrin (green), anti-pMad (red). **(P)** The statistical analysis of the average number of GSC and CB and Cysts in *eggpl*^*[1]*^ line. Error bars show SEM, ns indicates no significant difference, \*\*\**P*<0.001.

### 2.12. BrdU incorporation

5-Bromo-2’-deoxyuridine (BrdU), an analog of the nucleoside thymidine, was used in examining the frequency of the S-phase during cellular cycle in this study. The BrdU (Sigma-Aldrich, cat. no. B5002) saturated solution was freshly diluted with 200 μL PBS (10 mM) and 800 μL dehydrated alcohol. The female flies were dissected in petri dish containing 1 mL Schneider’s insect medium at room temperature, then the ovaries were carefully transferred into a 12-well plate with the mixture of BrdU solution and Schneider’s insect medium (1:100) for further incubation at 25°C. After 45 min, the ovaries were rinsed with Schneider’s insect medium for 2 times and washed with PBS 1 × 5 min respectively, and fixed in 4% PFA for 50 min. The samples were washed 2 × 10 min with 0.3% PBT (0.3% Triton X-100 in PBS), sequentially, washed with 0.6% PBT (0.6% Triton X-100 in PBS) for 45 min. Acid-treating the ovaries with 1 mL 0.6% PBT and 1mL 3.2 mol/L HCL for 30 min, and then the ovaries were washed with 0.3% PBT 3 × 10 min and 0.1% PBT for 30 min. The ovaries were blocked by 10% NGS (10% normal goat serum in 0.1% PBT) for 1 h and incubated with 1: 50 mouse anti-BrdU monoclonal antibody (Becton Dickinson cat. no. 7580) at 4°C, overnight. The next step was followed by common immunofluorescence staining procedure.

### 2.13. Protein overexpression of eggpl in Sf9 cell line in vitro

Briefly, the ORF of *eggpl* was inserted into vector piztv5-His (Novagen) in our laborary to overexpress eggpl, and cell co-transfection was carried out using FuGENE HD Transfection Reagent (Promega). Then, *Sf9* cell line was maintained at 28 °C in 25 cm^2^ culture flasks (Nest, China) in Grace’s insect cell culture medium containing 10% fetal bovine serum (Gibco, USA).

### 2.14. Statistical analysis

All data were analyzed by one-way ANOVA with *Duncan’s* multiple range test (DMRT) using a SAS statistical windows 8.1 package program (Microsoft, USA). *p*<0.05 was considered to be statistically significant.

## 3. Results

### 3.1. Overview of single cell RNA transcriptional atlas of Drosophila ovary

To characterize the transcriptional profile of ovarian cell types, we performed scRNA-seq on 7-day-old adult *Drosophila* ovaries by using 10× Genomics Chromium system to complete the complementary DNA (cDNA) synthesis and amplification, library preparation, and sequencing process (**Fig. 1A**). We then used t-Stochastic-Neighbor Embedding (t-SNE) in Seurat (24) to reduce the dimensionality and visualize the unsupervised cell distribution and 12 cell clusters (**Supplementary Fig. 1A**) was first classified based on their unique transcriptional profiles with the default resolution.

We further assigned cell types using canonical marker genes and further adjusted the Seurat resolution as needed, resulting in 24 distinct clusters in total (**Fig. 1 C and D**). The two clusters (cluster6 and cluster8) that expressed the germ cell marker *vasa* (25) were combined together as one germ cell cluster (**Supplementary Figure 1A**). We further divided the combined cluster into four subclusters (undifferentiated germ cell, oocyte, early nurse cell and later nurse cells) based on distinctions between the transcriptional profiles revealed by unsupervised clustering and identification of stage-specific markers (**Supplementary Fig. 1D**) (**Fig. 2A**). Both *nanos* which is maternally loaded into pole plasm and translated after fertilization (26) and *osk* which is highly enriched in germ plasm and accumulated in pole cells (27-28) were used to identify 3 subtypes of pole cell clusters at 0.5 resolution value (R=0.5). The stalk cell cluster was enriched in the expression of *zfh-1*(29), *stl* and *CG46339*, as expected (30). The cluster with upregulation of *Wnt4* and *GstS1*was considered as escort cell cluster according to the previous report (30). The polar cell cluster was identified by a known marker *upd1* (31) at default resolution value (R=1). Three subpopulations of stretch cells were distinguished by several marker genes, including *peb* (32), *sosie* (33), br (34), *past1, Glut4EF* and *Vha16-1* (35) at 0.5 resolution value (R=0.5), and the terminal follicle cell cluster was identified by the expression of *past1*. The expression of *SPARC* and *Jupiter* was used to identify the 1-4 stage of main body follicle cell (1-4 MBFC) (30). Four subclusters such as MBFC (stg. 9-10A) 1, MBFC (stg. 9-10A) 2, MBFC (stg. 10B) and cent/post MB 8 were identified by *psd, Vml, ttk, dec-1, bond* and *Fcp3C* (35) at 0.5 resolution value (R=0.5). The MBFC (stg. 12) and MBFC (stg. 14) were easily identified by maker genes such as *Femcoat, Ilp8* and *yellow-g* at 0.5 resolution value (R=0.5). The well-recognized marker genes *Hlm* and *su(r)* were used to identify hemocyte and plasmatocyte respectively (35, 36).

### 3.2. Identification of GSC cluster and 2 distinct undifferentiated germ cell subpopulations

To refine our previous clustering results, the GSC differentiation-related markers, such as *bam, bgcn, blanks* and *cycE* (30, 37), were used to identify undifferentiated germ cells, while *egg, put* and *cona*, which were known to be required in GSC maintenance and oogenesis (38-41), were used to identify the later stages. The enrichment of *c(3)G, Iswi* and *mael* was observed in the undifferentiated germ cell, early nurse cell and oocyte clusters, additionally, *CG15628* was observed specifically in the oocyte (**Fig. 2C**).

To characterize the spatial and temporal changes in transcription that occur during the initial stages of germ cell differentiation, we used Monocle3 to construct the developmental trajectory of the 4 germ cell clusters (**Supplementary Fig. 1C**). This analysis produces a graph-based trajectory called pseudotime that predicts the transcriptomic changes along the putative timing of developmental process (42). In this case, 4 germline clusters were arranged into a linear trajectory consisting of 3 branches, which is consistent with the continuous progression of germline development. We identified a subset of *bam*^*–*^ cells at one end of the trajectory that we concluded were GSCs (**Fig. 2B-B’**), and we assigned the remaining non-*bam* expressing cells and bam-positive cells to two distinct undifferentiated germ cell subclusters, namely undifferentiated germ cell-1 and undifferentiated germ cell-2. To further investigate the hypothesis and examine the putative GSC cluster, we plotted the GSC-related gene expression in pseudotime. Similar expression profiles of functional GSC differentiation genes (*bam, bgcn, out, twin* and *Set1*) (43-46) and GSC maintenance genes (*aret, Mei-P26, egg, Myc* and *Hrb98DE*) (47-51) were showed in trajectory (**Fig. 2D**). Consistent with our expectations, we observed low expression of genes associated with differentiation and high expression of genes associated with self-renewal, while their expression patterns along the trajectory were gradually changed over time. The expression patterns of top 5 expressed genes in three germline subclusters suggested that the developmental states of cells in GSC cluster and undifferentiated germ cell-1 cluster were similar to each other but different from that of the undifferentiated germ cell-2 cluster (**Fig. 2C**).

### 3.3. Construction of gene regulatory network in GSC

Although a large number of genes are expressed in all germ cells, some genes that are differentially expressed during early germ cell development may play more important role for GSC fate. To identify these types of genes, we conducted a comparative analysis on the transcriptional profile of 6 germ cell subclusters. We identified subsets of genes in the germline-1 and -2 clusters, early nurse cell cluster, oocyte cluster and later nurse cell cluster, that are differentially expressed compared to the GSC cluster (**Fig. 3A**). In addition, we performed an RNAi screen of 33 differentially upregulated genes in GSCs *vs* undifferentiated germline-1 and GSCs *vs* undifferentiated germline-2 clusters by using *nos-Gal4/CyO;tub-Gal80*^*ts*^*/TB*, a temperature-sensitive fly line, to individually trigger the available UAS-RNAi lines at adult stage. We found that RNAi knockdown of 19 upregulated genes induced disruption of GSCs/CBs homeostasis. Of these, 12 genes were classified as “changes to the number of GSC/CB”, 6 genes as “empty germarium” and 4 genes exhibited “differentiation defects” (**Fig. 3B-C**). Lastly, we scored the differentially expressed genes (score > 980) and constructed an interaction network (**Fig. 3D**). This analysis revealed a dense network of interactions between the differentially expressed genes, with genes that regulate translation (eIF2gamma, Ns1, and Prp19) forming major nodes. In addition, we found that RNAi knockdown of 21 out of 39 most highly expressed genes also caused a significant increase or decrease in the number of GSC/CB per germarium. (**Supplementary Fig. 2A-B**).

To identify the transcription factor-based gene regulatory network in different kinds of germ cells, we applied SCENIC analysis to our single-cell RNA sequencing data with 6 known germ cell types. The analysis revealed that the Dref, mal-f, Hsf, and REPTOR-BP regulons were enriched in GSC cluster, suggesting that they may play an important role in the regulation of early GSC development (**Fig. 3E**). GO and KEGG enrichment analysis provided additional information about the biological processes that are enriched during germ cell development. Specifically, we found that the enriched GO terms were closely related to the basic physiology of *Drosophila* such as cellular metabolic process, protein catabolic process, cytoskeleton organization and cell cycle. Notably, a proportion of functional GO terms in GSC cluster was particularly enriched in ubiquitin-dependent protein catabolic process and cellular catabolic process, which was in relation to cancer and disorder research (52-53) (**Supplementary Fig. 2C**). The top 10 pathways in the KEGG enrichment analysis revealed that the differentially expressed genes in GSC, undifferentiated germ cell-1 and -2 were significantly enriched for DNA replication and disease related pathways, while early nurse cell, oocyte and later nurse cell specifically enriched in the pathway of ribosome, Hippo signaling pathway and MAPK signaling pathway (**Supplementary Fig. 2D**).

### 3.4. Validation of candidate markers genes in germ cell

To identify new markers of distinct stages of germ cell differentiation, we selected 10 candidate genes that are predicted to be expressed in GSCs by pseudotime analysis and assayed their expression patterns by *in situ hybridization*. These included 9 GSC specific markers and 1 germline cysts marker (**Fig. 4A**). We identified seven genes that were specifically expressed in the anterior tip of the germarium, where the GSCs are located, including one gene, *CG32814*, which we named *eggplant* (*eggpl*) because knockdown causes an enlarged ovary with many retained eggs, as described below (**Fig. 4B**). To validate these expression patterns at the protein level, we generated antibodies against two genes, the basal transcription factor TfIIA-S (54) and Eggpl. Indeed, we found that the immunofluorescence signals of both antibodies were highly enriched in GSC and early germ cells, consistent with our in situ hybridization results (**Fig. 4C, E**). TfIIA-S was localized to the nucleus, as expected for a transcription factor, whereas the Eggpl was enriched in the cytoplasm, which we confirmed *in vitro* using the *Sf9* cell line (**Fig. 4F**). In addition to these genes with highly specific expression patterns, we also found that *hang*, a conserved regulator of ethanol tolerance (55), is expressed in germ cells throughout the germarium, and that *CG7255* is expressed in germ cell cysts and nurse cells but not in GSCs. These expression patterns also align with the order of expression of *eggpl, hang* and *CG7255* predicted by pseudotime analysis (**Fig. 4D**).

### 3.5. The unique expression patterns of eggpl in germline

To further characterize the cells that express *eggpl*, we co-labeled for *eggpl* mRNA and either α-Spectrin protein, which localizes to a cytoplasmic structure that is spherical in GSCs (called “spectrosomes’) and elongates to “fusomes” in cystoblasts **(56)**, or the germ cell-specific protein, Vasa (**Fig. 5A-B**). We found that *eggpl* mRNA was specifically enriched in GSCs. To confirm this observation, we probed for *eggpl* in a *bam-GFP* line. Indeed, we found that *eggpl* transcript was enriched in the Bam-GFP^−^ cells at the anterior tip of the germarium, consistent with GSC-specific expression (**Fig. 5D**).

To assess the pattern of Eggpl protein expression, we constructed and eggpl::GFP line in which GFP was knocked into the endogenous locus (**Supplementary Fig. 3 A**). We co-stained for GFP and α-Spectrin, pMad (**Fig. 5E**) or anti-Bam (**Fig. 5F**) and found that, in contrast to the *eggpl* mRNA expression pattern, Eggpl::GFP was detectable in germ cells throughout Region 1 (**Fig. 5E**). Interestingly, Eggpl::GFP protein levels varied by stage, with the highest level of expression in the cells just downstream from the GSC niche (**Fig. 5L-M**). Thus, the range of protein expression was broader than the range of mRNA expression and protein levels and was actually highest in cells that have no detectable *eggpl* transcript. In addition, we constructed a *UAS-eggpl-GFP* line and examined the overexpression pattern of *eggpl* by using mRNA *in situ hybridization* (**Fig. 5G and 5I**). The result showed broad expression of *eggpl* in Region 1 of the overexpression line. Together, these results suggest that there are distinct layers of regulation of *eggpl* gene expression at the mRNA and protein levels.

### 3.6. Ectopic Expression of eggpl affects the differentiation of Germline Stem Cells and Cystblasts

To determine whether *eggpl* is involved in the regulation of GSC fate, we expressed *UAS-eggpl-RNAi* and *UAS-eggpl-GFP* lines with *nos-Gal4;tub-Gal80*^*ts*^ line when the adult flies emerged from pupae, and stained for pMad and α-Spectrin to identify GSC and early germ cells (**Fig. 5H**). With this combination of markers, GSCs were identified as cells at the tip of the germarium that have high levels of pMad and spherical α-Spectrin^+^ spectrosomes whereas the differentiating germline cysts (2-, 4-, 8- and 16-cell) have low levels of pMad and α-Spectrin^+^ interconnecting branched fusomes (56). We did not observe a significant difference in GSC or CB number upon knockdown of *eggpl* (**Fig. 5J**) but found an increase in branched cysts (2-cell and 4-cell stages) upon knockdown of *eggpl* (**Fig. 5K**). To further study *eggpl* function, we generated an *eggpl* allele, *eggpl*^*[1]*^, using CRISPR. Eggpl protein was undetectable in *eggpl*^*[1]*^ germaria (**Fig. 5N**), indicating that the allele disrupts protein expression. Consistent with our RNAi results, both the number of GSC and the number of CB were not affected (**Fig. 6E-F**) in *eggpl*^*[1]*^, while the number of germline cysts in the 2-cell to 8-cell stages was significantly increased (**Fig. 5O-P**). Taken together, these results indicate that *eggpl* is required for GSC differentiation in *Drosophila* ovary.

**Fig. 6.**
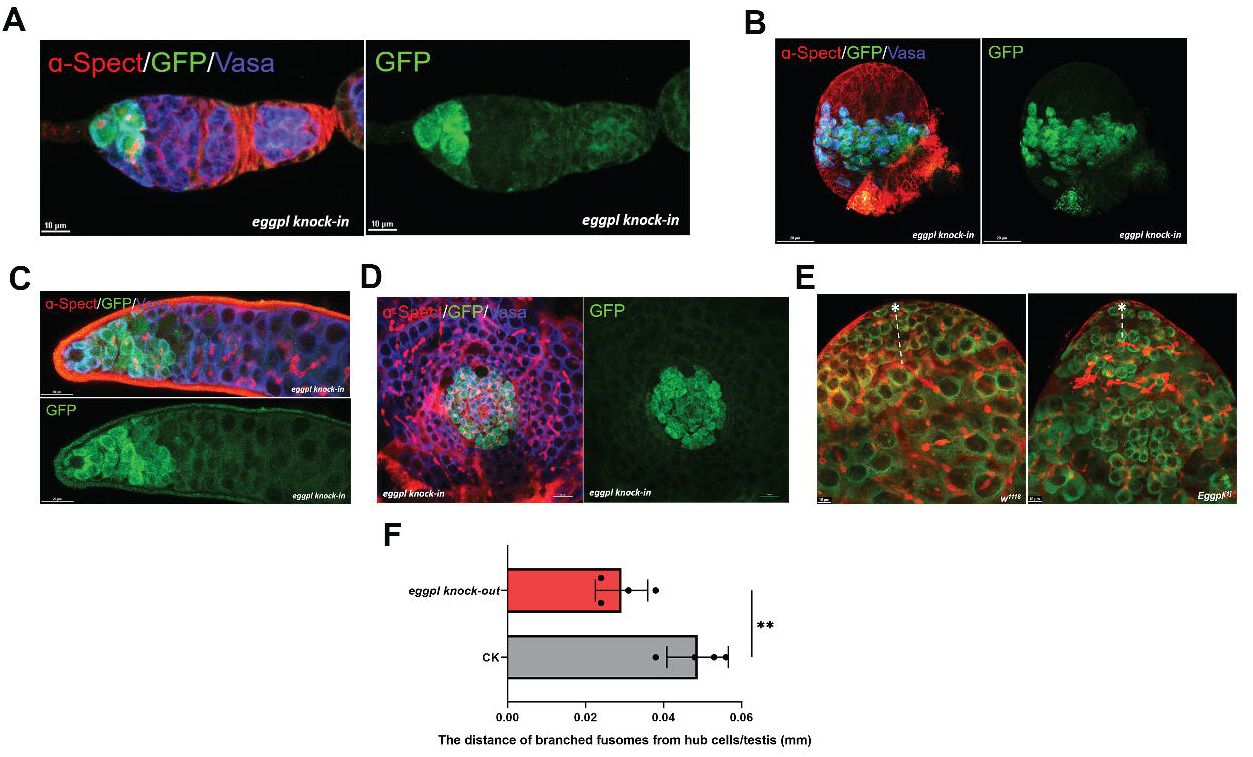
The dynamical expression of *eggpl* in testis and ovary. **(A)** Immunofluorescence staining with anti-α-Spectrin (red), anti-Vasa (blue) and anti-GFP (green) on adult ovary. **(B)** Immunofluorescence staining on PGCs of larvae ovary. **(C)** Immunofluorescence staining on GSCs and early spermatogonia in adult testis. **(D)** Immunofluorescence staining on PGCs of larvae testis tissue. **(E)** Immunofluorescence staining with anti-α-Spectrin (red) and anti-Vasa (green) on testis from wild type or *eggpl*^*[1]*^ flies. **(F)** The measurement of distance between hub cells and early germ cyst. n=4, error bars show SEM, \*\**P*<0.01.

### 3.7. The expression of eggpl in GSCs and primordial germ cells (PGCs) in both ovary and testis at different developmental states

Many genes regulate germ cell differentiation in both males and females, and, indeed, we found that *eggpl* is also expressed in male GSCs and early spermatogonia (**Fig. 6C**). Since the *Drosophila* GSCs are derived from a small population of primordial germ cells (PGCs) with undifferentiated states, the profiles of gene expression in PGCs may vary from that of the adult. To detect whether the expression of *eggpl* may be more widely exhibited in germline lineage from larvae to adult, we dissected the gonads from male and female larvae in *eggpl knock-in* lines, and stained with anti-α-Spectrin and anti-Vasa to label the PGCs and germ cells. We found that *eggpl* is expressed in both male and female larvae PGCs and early undifferentiated germ cells (**Fig. 6B and 6D**), suggesting that *eggpl* may function at these early stages as well. In the testes of wild type male flies, GSC were present next to the apical tip of testes and gradually differentiated into spermatogonial cells with germline specific branched organelle fusomes. While we found that the distance of branched fusomes from the hub cells in *eggpl knock-out* testes is significantly (*p*<0.01) less as compared to those in control (**Fig. 6E-F**). This finding suggested an early onset of premature differentiation of GSCs in the testes when *eggpl* was lost.

### 3.8. Depression of eggpl increases egg production and regulates germ cell proliferation

Since the disruption of *eggpl* led to an increase in the frequency of germ cell cysts, we examined the oviposition on *eggpl-RNAi* and *eggpl*^*[1]*^ lines, and found a significant increase in the number of eggs laid by flies with RNAi knockdown of *eggpl* in germ cells compared to sibling controls. (**Fig. 7B**). In addition, we noticed that the size of the whole ovary and the number of mature eggs per ovary were substantially increased upon RNAi and knock out of *eggpl* in germ cells at 2-, 7-, and 14-day-old flies but returned to a size that is comparable to wildtype by 21-days (**Supplementary Fig. 3 B**). Based on this phenotype, we named the gene *eggplant*. Furthermore, we surmised that *eggpl* could be involved in the regulation of proliferation. To test this possibility, we assayed for proliferation in germ cells using a BrdU incorporation assay, which identifies cells in S-phase (**Fig. 7A**). Compared with wild type fed on standard diet, the average number of BrdU^+^ cysts was significantly increased in germ cell cysts in wild type fed on rich yeast diet, *eggpl-RNAi* (on standard or Rich diet), and *eggpl*^*[1]*^ (on standard or Rich diet) lines respectively (**Fig. 7C**).

**Fig. 7.**
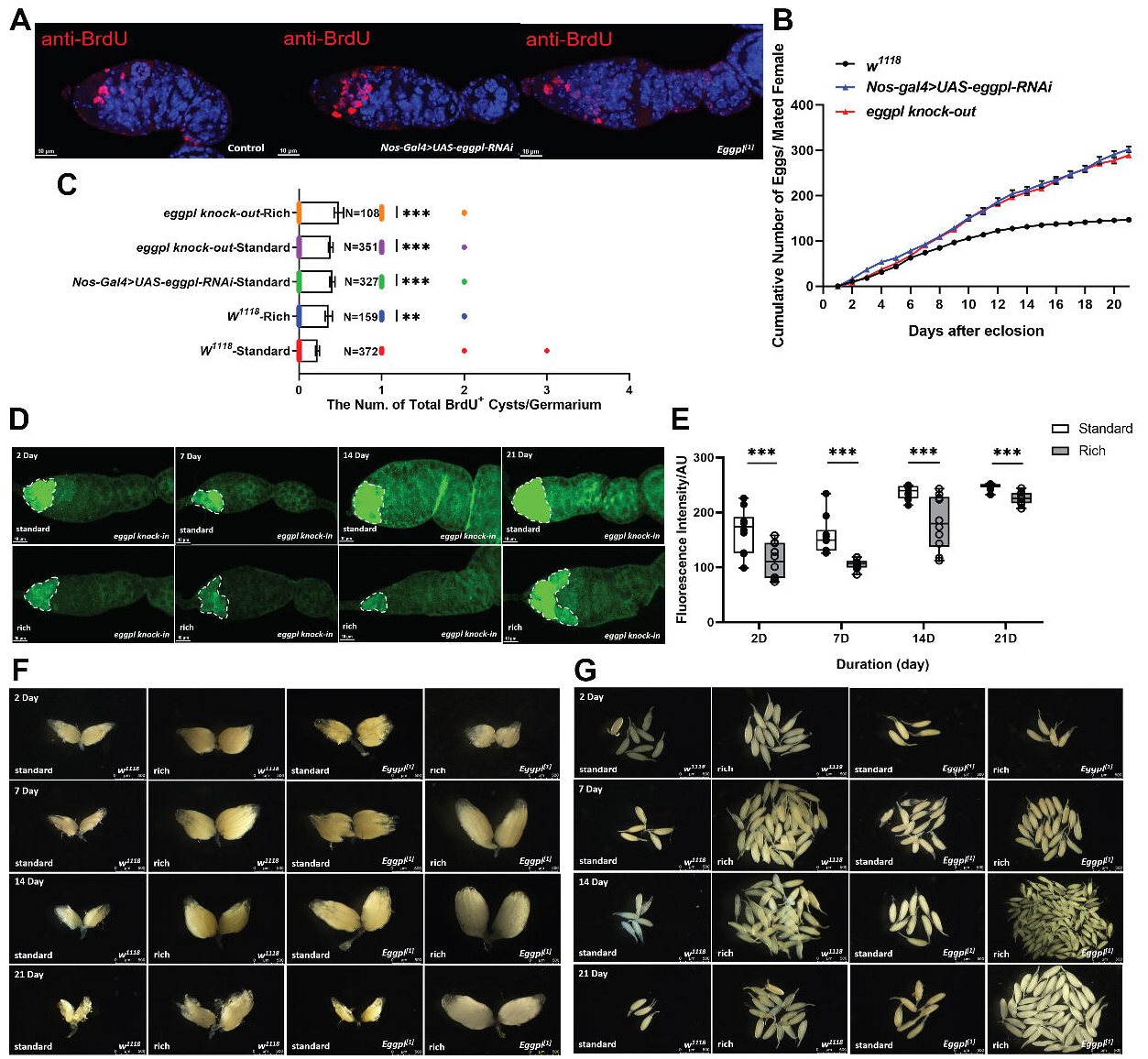
The function of *eggpl* might be involved in regulating cell cycle of germline cysts. **(A)** Immunofluorescence staining with anti-BrdU (red). **(B)** The oviposition of wild type, *Nos-Gal4*>*UAS-eggpl-RNAi* and *eggpl*^*[1]*^ line over 21 days. **(C)** The number of BrdU^+^ germ cell cysts. Error bars show SEM, ns indicates no significant difference, \*\**P*<0.01, \*\*\**P*<0.001. **(D)** Immunofluorescence staining with anti-GFP on germaria from *eggpl::GFP knock-in* flies with or without feeding fresh yeast paste on day 2-, 7-, 14- and 21. The dotted line indicates the examined area. **(E)** Quantification of the GFP intensity mean in the *eggpl::GFP knock-in* line, n=10, \*\*\**P*<0.001. **(F-G)** Images of ovaries (F) or ovarioles (G) from flies of the indicated genotypes, feeding conditions, and days post-eclosion showing comparisons of the overall ovary sizes (F) and the numbers of retained matured eggs (G).

The increase in ovary size and GSC proliferation that we observed upon knockdown of *eggpl* phenocopies the response of ovaries to a rich protein diet, (i.e. daily feeding of wet yeast paste). This suggests that the wet yeast diet may be promoting oogenesis in part by repressing *eggpl* expression. To test this hypothesis, we compared the ovaries from control and *eggpl*^*[1]*^ that were maintained on standard food or standard food plus wet yeast paste. The ovary size of the control flies was substantially increased by the addition of wet yeast paste, consistent with previous reports **(16)**. Interestingly, we found that the ovary size and number of eggs in *eggpl* knockout lines maintained on either standard food alone or on standard food with wet yeast were comparable to the controls that were maintained with wet yeast paste (**Fig. 7F-G**). In addition, we found that the intensity of *eggpl* signal was significantly decreased in flies that were maintained with wet yeast paste (**Fig. 7D-E**). Wet yeast paste in the diet is known to promote increased egg production by signaling to the GSC niche through the insulin pathway. Therefore, taken together, these observations suggest a model in which a high yeast diet promotes GSC proliferation and increased egg production by inhibiting the expression of *eggpl* in GSCs and early germ cells, perhaps downstream of insulin signaling.

### 3.9. The eggpl mediates GSC differentiation via the MMP-dependent Timp regulation

The extracellular matrix (ECM) is an important remodeling component of ovarian niche, which is responsible for the cellular organization, cell-matrix adhesion and tissue stiffness (57). It is composed of Laminins, Perlecan, Collagen IV, Glutatin and mucin-type O-glycoproteins (58-59), and the ECM composition is regulated by a family of proteolytic enzymes, matrix metalloproteinases (MMPs) (60). Tissue inhibitors of metalloproteinases (TIMPs) mediate the inhibition of MMP activity, which is accomplished by blocking the MMPs catalytic domain with the amino and carbonyl groups of the TIMP N-terminal cysteine residue (61). We hypothesized that it may be regulated by MMP-dependent *Timp* signaling in the early GSC lineage. To test this hypothesis, we assayed for changes in Eggpl protein levels upon overexpression and RNAi knockdown of MMP-Timp related genes. We found that the fluorescence intensity of Eggpl protein levels were decreased upon *Timp* knockdown, and increased when *Timp* was overexpressed. In addition, overexpression of either *Mmp1* or *Mmp2* decreased the fluorescence intensity of Eggpl protein level. Conversely, Eggpl protein levels were slightly increased upon *Mmp1* RNAi (**Fig. 8A**). Interestingly, RNAi knockdown of *Timp* and *Mmp1/2* overexpression caused an enlarged ovary phenotype, similar to the phenotype we observed upon RNAi knockdown of *eggpl*, while *Timp* overexpression and RNAi knockdown of *Mmp1* did not affect ovary size (**Fig. 8B-C**). Collectively, these results support a model in which *eggpl* mediates the GSC differentiation process *via* MMP-dependent Timp regulation pathway (**Fig. 8D**).

**Fig. 8.**
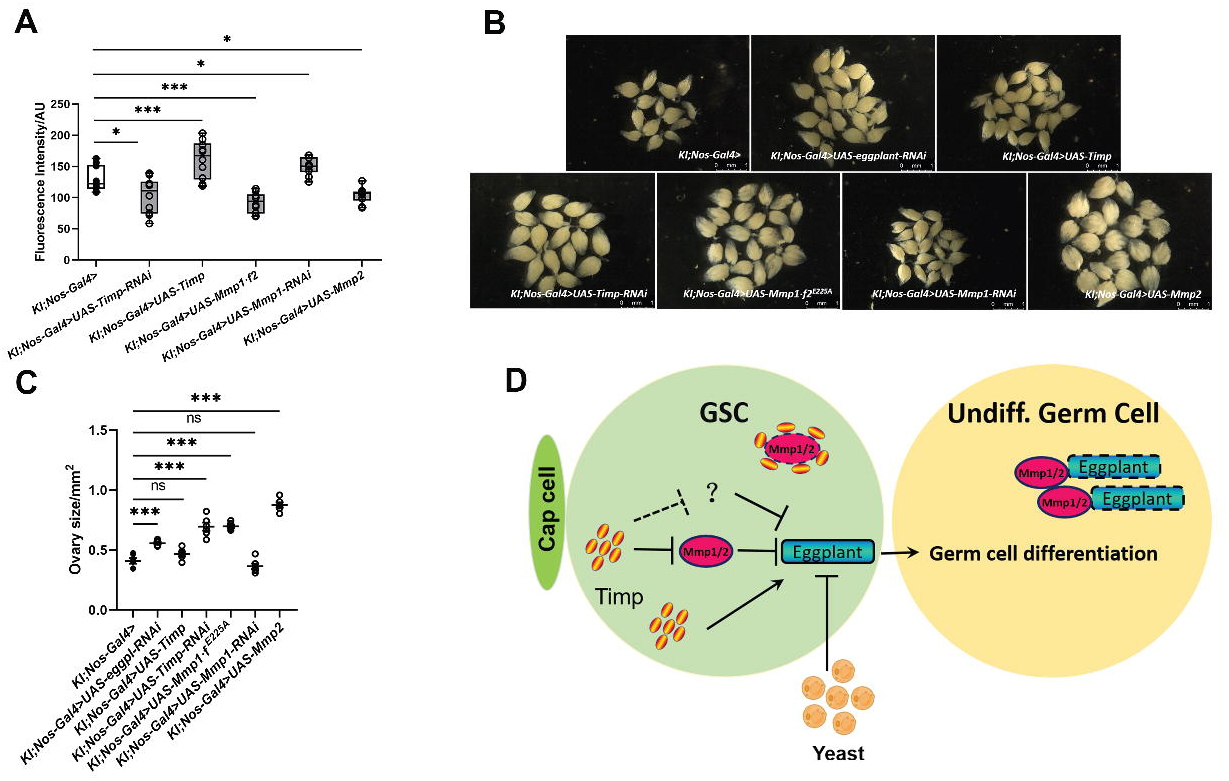
The *eggpl* maintains GSCs differentiation *via* MMP-dependent Timp regulation. **(A)** Quantification of the GFP signal in germaria from *eggpl::GFP knock-in* flies of the indicated genotypes. n=10, \**P*<0.05, \*\**P*<0.01, \*\*\**P*<0.001. **(B-C)** Images of ovaries (B) and quantification of ovary size (C) from flies of the indicated genotypes. Error bars show SEM, ns indicates no significant difference, \*\*\**P*<0.001. **(D)** Model depecting the regulation of *eggpl* expression and its role in GSCs and differentiating cysts.

## 4. Discussion

Decades of research have established the *Drosophila* female germline stem cell as a favorable system to understand germ cell and stem cell biology. The GSCs are required in germarium to support continuous production of differentiating germ cells throughout most of adulthood. The GSCs are typically identified by their localization at the anterior tip of the germarium (62), and the presence of spectrosomes or high pMad-signal (63), and many other useful marker genes have also been described. For example, the expression of *Lamin C*, a typical marker, is strongly expressed in TF cells, displays the weak expression in cap cells, and is not detectable in escort cells. Conversely, *traffic jam* (*tj*) is highly expressed in escort cell and cap cells, but not detectable in TF cells (64). These markers facilitate many tissue- and cell type-specific genetic manipulations *in vivo*, which can be used to understand the gene functions and signaling pathways that regulate germ cell differentiation. Therefore, discovery of specific marker genes in GSCs will have many applications in the study of germ cell and stem cell biology. However, it has been difficult to identify new markers of GSCs, in part because they are rare in wildtype tissue, and thus not amenable to bulk sequencing approaches. In our study, we performed 10×single cell transcriptomes sequencing on whole ovary, and identified 24 distinct cell populations by using known marker genes (**Fig. 1C-D**). Taking into account the variation of the sample, we compared our data with 4 public *Drosophila* ovary scRNA-seq datasets (30, 65). The plotting results showed that our dataset is well integrated with others (**Supplementary Fig. 1 B**). Here, we analyzed the 175 cells in the GSC cluster to identify individual genes and gene regulatory networks that may be important for GSC function. This approach produced a list of genes that is highly enriched for genes that produce a phenotype when knocked down in germ cells by RNAi or that have a specific expression pattern in the early GSC lineage. This validates the approach and provides a new resource for the community.

The evolutionarily conserved insulin-like growth factor (IGF) pathway has multiple roles in the modulation of GSC proliferation and maintenance. A protein-rich diet induces the production of insulin-like peptides (DILPs) in the brain, which regulate GSC division and cyst growth on a protein-rich diet (14). Several intracellular signals have been identified that function downstream of insulin signaling in germ cells. These include phosphoinositide-3 kinase (PI3K), *dFOXO*, and cell cycle factors, such as CycA, CycB, CycE and E2F1 (66, 67). However, little is known about the gene targets of this pathway that modulate the rate of differentiation. Our findings that a rich yeast diet causes a decrease in *eggpl* expression and that knockdown or knockout of *eggpl* mimics the effects of a rich yeast diet on the ovary raises the interesting possibility that *eggpl* may be a key link between nutritional cues and the regulation of oogenesis (**Fig. 7C-G**). It is interesting that knockdown or knockout of *eggpl* is sufficient to induce such a substantial increase in egg laying on standard food without yeast supplementation. This suggests that protein in the diet is not the limiting factor under these conditions.

As the endogenous inhibitors of MMPs activities, TIMPs have been reported to regulate a series of cellular processes including neurite differentiation, apoptosis and cell division (68-70). Increasing evidence indicates that Timp-mediated inhibition of MMP activity in the extracellular matrix could reduce hepatocyte proliferation in a murine regeneration model **(71)**. In *Drosophila*, the regulation of MMPs (*Mmp1* and *Mmp2*) activity by inhibitory TIMP plays a key role in tissue stiffness and ovarian niche organization. For instance, *timp* regulates the distribution of *Mmp1* and *Mmp2*, which could maintain GSC niche homeostasis and interfollicular stalk formation. The *loss of timp* causes the defects on organization of germline cysts **(57)**. Recently, it has been shown that mRNA expression of *timp* strongly enriched in the place where GSCs reside, and the Mmp1 and Mmp2 protein accumulated in the most anterior of germarium (57, 72). Our results suggest that Eggpl in early germ cell may be mediated by MMP-dependent Timp pathway (**Fig. 8A**). Further investigation will be needed to explore how TIMP dependent inhibition of MMP regulates GSC division and whether TIMP functions independent of MMP inhibition in germ cells.

Taken together, our research aimed to unveil the developmental features of GSCs. The bioinformatics analysis allowed us to obtain the transcriptomes of 175 GSCs, and provided a transcriptional perspective of two distinct undifferentiated germ cell clusters. The novel GSCs marker genes validated in this study were beneficial to better understand the signature of stem cell lineage. We further introduced a GSC specific functional gene, *eggpl*, and explored its gene function in GSC differentiation progress. On the other hand, the combination of differentially expression gene analysis and RNAi screen allowed us to gain a better understanding of the potential genetic interactions between genes involved in GSC maintenance and differentiation.

## Acknowledgements

This work was supported by National Natural Science Foundation of China (No. 31572335). T.G.N is supported by a grant from the National Institutes of Health (GM136348). We gratefully thank Guangzhou Genedenovo Biotechnology Co., Ltd for the assistance in sequencing and bioinformatics analysis. Specially thank for the generous help and professional guidance provided by Suning Liu (South China Normal University, China).

## Figure legends

**Supplementary Fig. 1.**
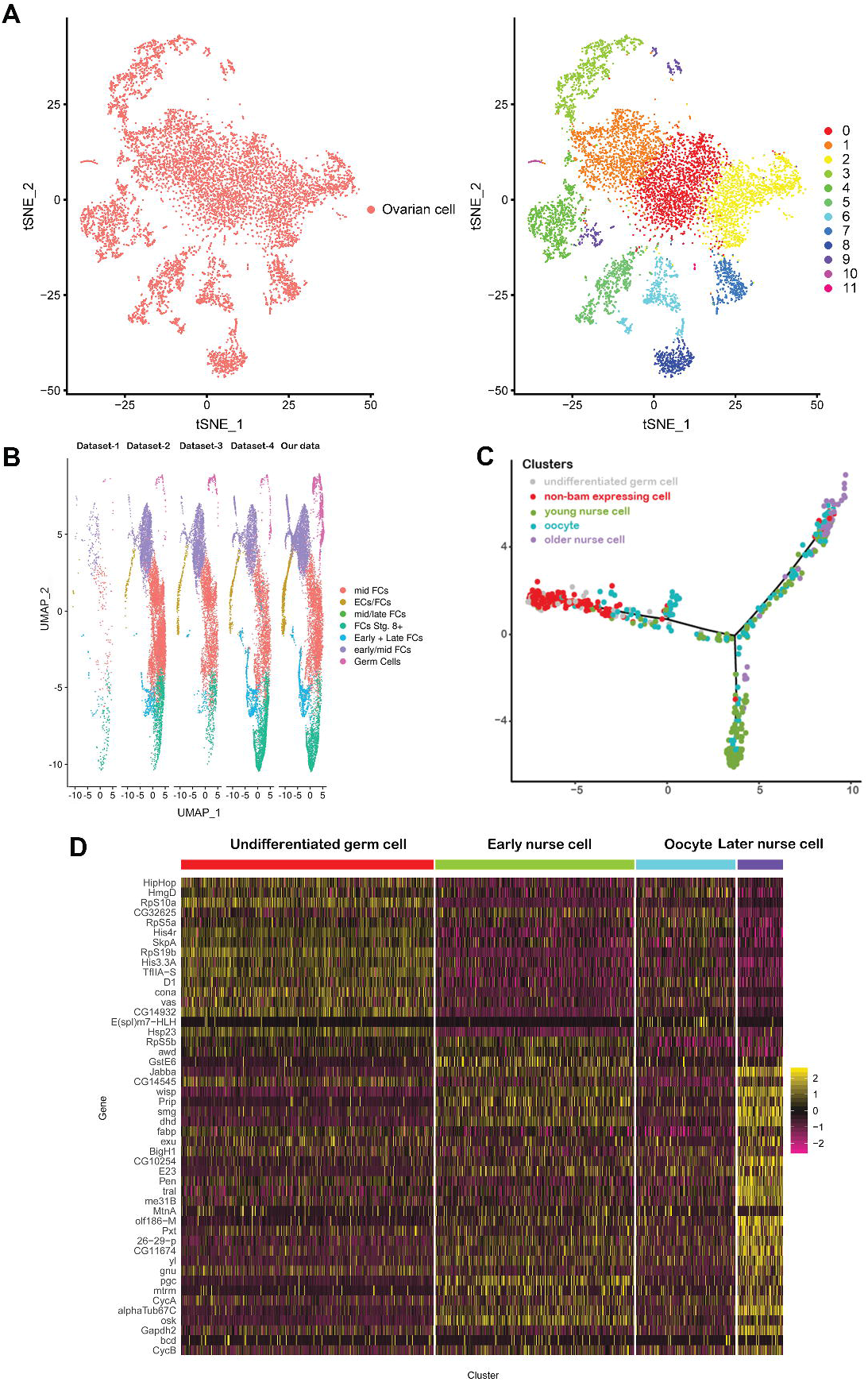
The preliminary analysis on scRNA-seq data. **(A)** The original tSNE plot showing the distribution of distinct clusters in our dataset. **(B)** UMAP plots showing a comparison of four published datasets with the dataset produced in this study. Dataset-1, dataset-2 and dataset-3 are from Rust, *et al*. 2020, Dataset-4 is from Jevitt, *et al*, 2020. **(C)** The trajectory of undifferentiated germ cell, non-bam expressing cell, young nurse cell, oocyte and older nurse cell clusters in pseudotime, which contained *tj*-expressing cells and non-*vasa*-expressing cells. **(D)** The heat map showing the expression of known typical marker genes in each cluster.

**Supplementary Fig. 2.**
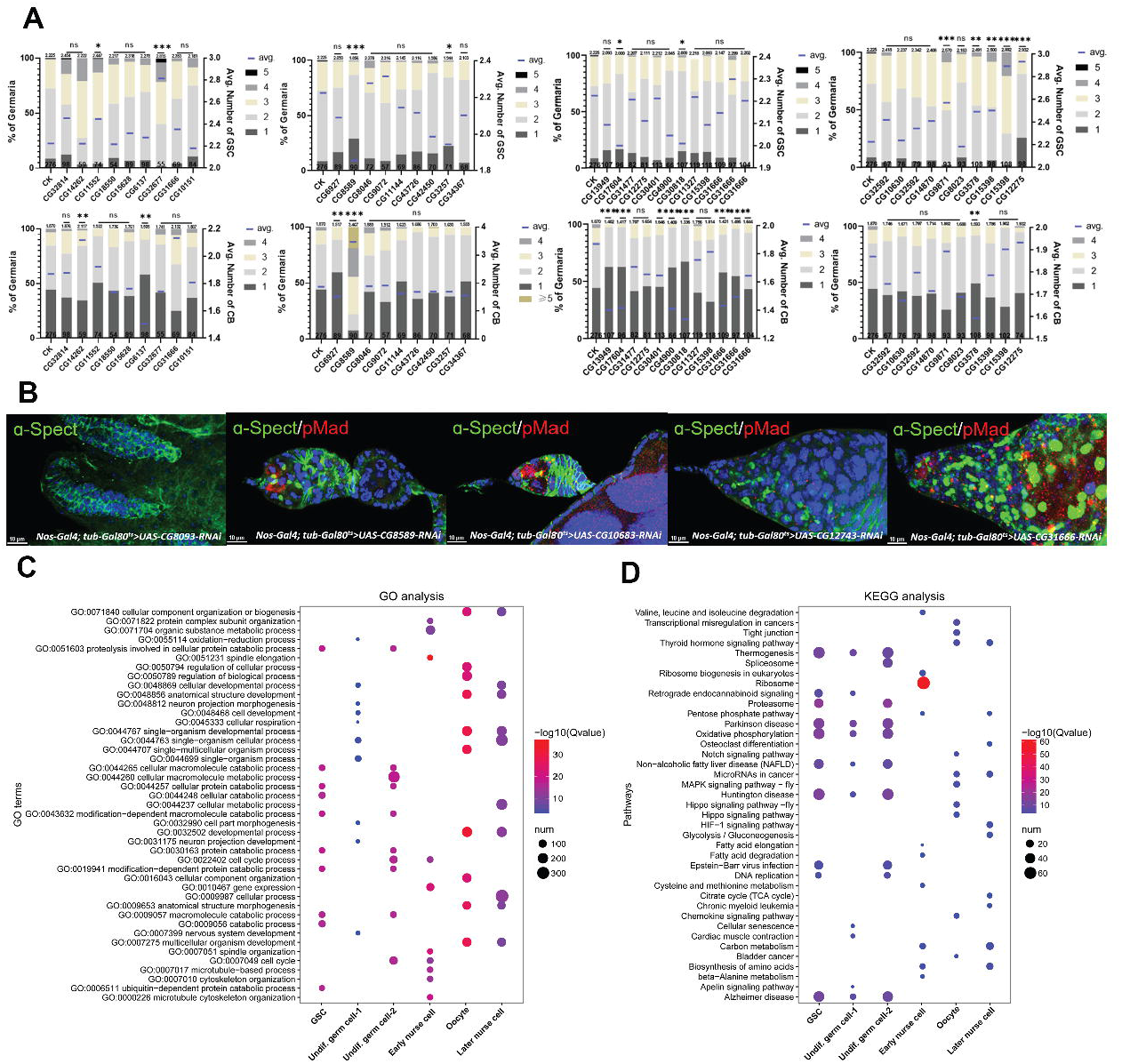
The RNAi screen on top expressed genes and enrichment analysis on GSCs. **(A)** The average number of GSCs and CBs in RNAi lines of screened top genes. Error bars show SEM, ns indicates no significant difference, \**P*<0.05, \*\**P*<0.01, \*\*\**P*<0.001. **(B)** The typical phenotypes of germaria in selected RNAi lines stained with anti-α-Spectrin (green) and anti-pMad (red). **(C)** Dot plot showing the enrichment of GO terms associated with the differentially expressed genes in the 6 germ cell subclusters. **(D)** Dot plot showing the KEGG enrichment results.

**Supplementary Fig. 3.**
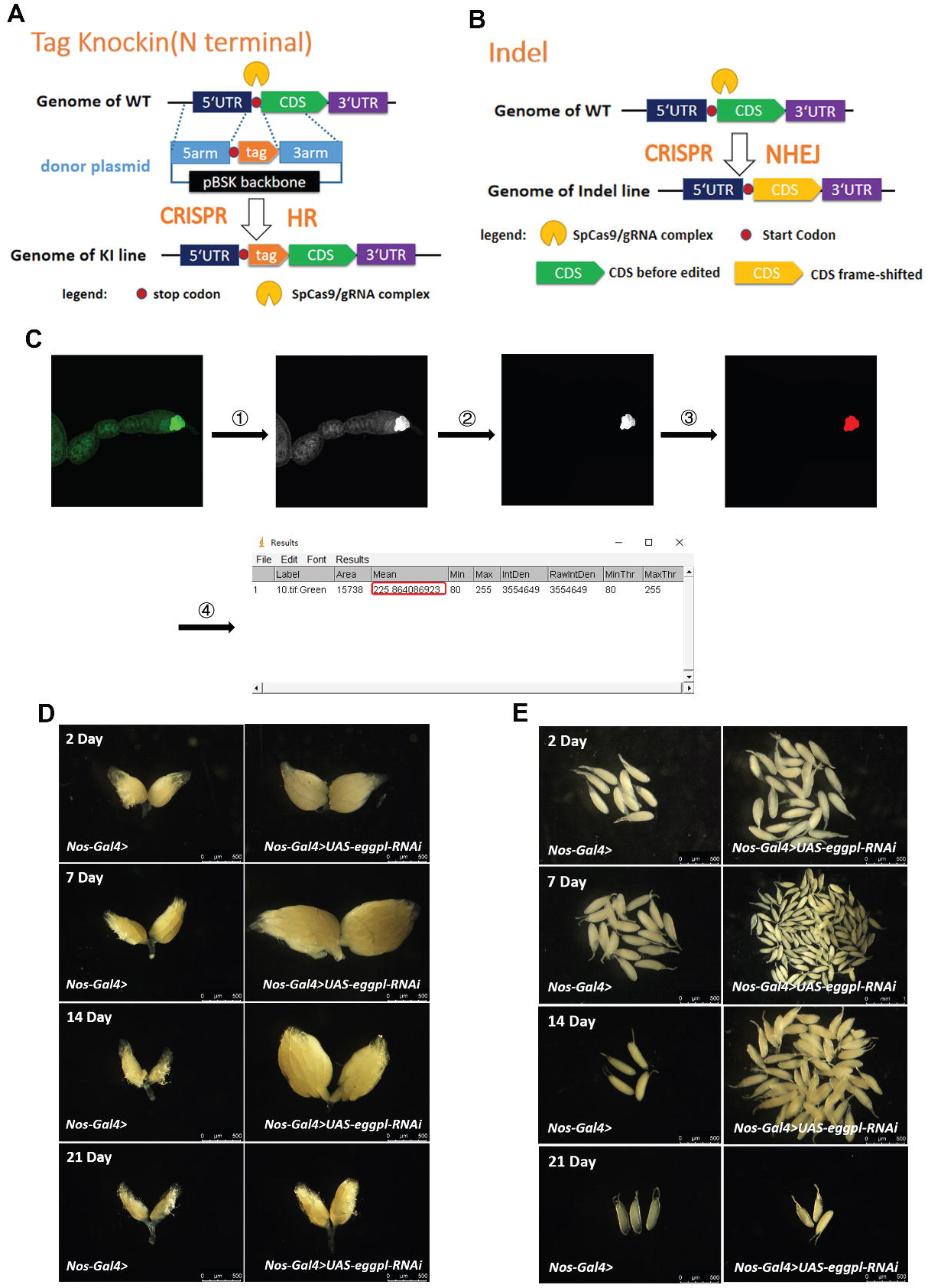
The schematic diagram of *eggpl knock-in* and *knock-out* lines construction and the ovary phenotype of *Nos*-*Gal4*> and *Nos*-*Gal4*>*UAS-eggpl-RNAi* Line. **(A)** The tag with GFP and 6HA was inserted on *N* terminal in fly genome. The CDS indicates coding DNA sequence of *eggpl*. **(B)** The construct of *eggpl*^*[1]*^ line was conducted by shifting the CDS frame. **(C)** The flow of immunofluorescence intensity analysis. The intensity values were analyzed with these command: 1. *Image > Type > RGB Stack*. 2. *Freehand selections > Cut*. 3. *Adjust > Threshold*. 4. *Analyze > Set Measurements > Area & Min & max gray value & Limit to threshold > Measure*. **(D-E)** The phenotypes of *Nos-Gal4* and *Nos*-*Gal4*>*UAS-eggpl-RNAi* ovary size and retained eggs on 2-, 7-, 14- and 21-day.

## References

(1) Flores H A, Dumont V, Fatoo A, et al. Adaptive Evolution of Genes Involved in the Regulation of Germline Stem Cells in Drosophila melanogaster and D. simulans[J]. G3-Genes Genomes Genetics, 2015, 5(4).

(2) Yoshinari Y, Ameku T, Shu K, et al. Neuronal Octopamine Signaling Regulates Mating-induced Germline Stem Cell Increase in Female Drosophila melanogaster[J]. eLife Sciences, 2020, 9:e57101.

(3) Merkle J A, Hinnant T D. Coordinating Proliferation, Polarity, and Cell Fate in the Drosophila Female Germline[J]. Frontiers in Cell and Developmental Biology, 2020, 8.

(4) Dansereau D A, Lasko P. The Development of Germline Stem Cells in Drosophila[J]. Humana Press, 2008.

(5) Tomotsune A, Ryusuke N, Liliane S. Mating-Induced Increase in Germline Stem Cells via the Neuroendocrine System in Female Drosophila[J]. PLOS Genetics, 2016, 12(6):e1006123..

(6) Wang L, Cai L Y. The JAK/STAT Pathway Positively Regulates DPP Signaling in the Drosophila Germline Stem Cell Niche[J]. Journal of Cell Biology, 2008, 180(4):721–728.

(7) Drummond-Barbosa D. Local and Physiological Control of Germline Stem Cell Lineages in Drosophila melanogaster[J]. Genetics, 2019, 213.

(8) Schulz C, Wood C G, Jones D L, et al. Signaling from Germ Cells Mediated by the Rhomboid Homolog Stet Organizes Encapsulation by Somatic Support Cells[J]. Development, 2002, 129(19):4523–4534.

(9) Jin S, Wei H M, Xu J, et al. Histone H1-mediated Epigenetic Regulation Controls Germline Stem Cell Self-renewal by Modulating H4K16 Acetylation[J]. Nature Communications. 2015, 6:8856

(10) Buszczak M, Paterno S, Spradling A C. Drosophila Stem Cells Share A Common Requirement for the Histone H2B Ubiquitin Protease Scrawny[J]. Science, 2009, 323(5911).

(11) Wang X, Pan L, Wang S, et al. Histone H3K9 Trimethylase Eggless Controls Germline Stem Cell Maintenance and Differentiation[J]. PLoS Genetics, 2011, 7(12):e1002426.

(12) Maines J Z, Park J K, Williams M, et al. Stonewalling Drosophila Stem Cell Differentiation by Epigenetic Controls[J]. Development, 2007, 134(8):1471–1479.

(13) Li Y, Zhang Q, Carreira-Rosario A, et al. Mei-P26 Cooperates with Bam, Bgcn and Sxl to Promote Early Germline Development in the Drosophila Ovary[J]. Plos One, 2013, 8(3), p. e58301.

(14) LaFever Leesa, Drummond-Barbosa, et al. Direct Control of Germline Stem Cell Division and Cyst Growth by Neural Insulin in Drosophila[J]. Science, 2005.

(15) Hsu H J, Lafever L, Drummond-Barbosa D. Diet Controls Normal and Tumorous Germline Stem Cells via Insulin-Dependent and -Independent Mechanisms in Drosophila[J]. Developmental Biology, 2008, 313(2):700–712.

(16) Drummond-Barbosa D, Spradling A C. Stem Cells and Their Progeny Respond to Nutritional Changes during Drosophila Oogenesis.[J]. Developmental Biology, 2001, 231(1):265-278(17)

(17) Su Y H, Rastegri E, Kao S H, et al. Diet regulates Membrane Extension and Survival of Niche Escort Cells for Germline Homeostasis via Insulin Signaling[J]. Development, 2018, 145(7):dev.159186.

(18) Hsu H J, Drummond-Barbosa D. Insulin Levels Control Female Germline Stem Cell Maintenance via the Niche in Drosophila[J]. Proceedings of the National Academy of Sciences of the United States of America, 2009, 106(4):1117–1121.

(20) Y Fu, X Huang, P Zhang, et al. Single-cell RNA Sequencing Identifies Novel Cell Types in Drosophila Blood[J]. Journal of Genetics and Genomics, 2020, 47(4).

(21) Avila Cobos, F., Vandesompele, J., Mestdagh, P. and De Preter, K.,. Computational Deconvolution of Transcriptomics Data from Mixed Cell Populations. Bioinformatics, 2018, 34(11), pp. 1969–1979.

(23) Davie K, Janssens J, Koldere D, et al. A Single-cell Transcriptome Atlas of the Aging Drosophila Brain[J]. Cell, 2018, 174(4):1–17.

(19) Hao Y, Hao S, Andersen-Nissen E, et al. Integrated Analysis of Multimodal Single-Cell Data. Cold Spring Harbor Laboratory, 2020.

(20) Laurens V D M, Hinton G. Visualizing Data using t-SNE[J]. Journal of Machine Learning Research, 2008, 9(2605):2579–2605.

(21) Labun K, Montague T G, Krause M, et al. CHOPCHOP v3: Expanding the CRISPR Web Toolbox beyond Genome Editing[J]. Nucleic Acids Research, 2019, 47(W1)

(22) Maurice L, Adams F F, Michelle N, et al. Refined sgRNA Efficacy Prediction Improves Large- and Small-scale CRISPR–Cas9 Applications[J]. Nucleic Acids Research, 2018(3):3.

(23) Bassett A, Tibbit C, Ponting C, et al. Highly Efficient Targeted Mutagenesis of Drosophila with the CRISPR/Cas9 System[J]. Cell Reports, 2014, 6(6):1178–1179.

(24) Satija R, Farrell J A, Gennert D, et al. Spatial Reconstruction of Single-cell Gene Expression Data[J]. Nature Biotechnology, 2015, 33(5):495–502.

(25) P F, Lasko Ashburner. The Product of the Drosophila Gene vasa is Very Ssimilar to Eukaryotic Initiation Factor-4A.[J]. Nature, 1988, 335:611–617.

(26) Deshpande G, Calhoun G, Yanowitz J L, et al. Novel Functions of nanos in Downregulating Mitosis and Transcription during the Development of the Drosophila Germline[J]. Cell, 1999, 99(3):0–281.

(27) Kim-Ha J, Smith J L, Macdonald P M. oskar mRNA is Localized to the Posterior Pole of the Drosophila Oocyte[J]. Cell, 1991, 66(1):23–35.

(28) Eichler C E, Hakes A C, Hull B, et al. Compartmentalized oskar Degradation in the Germ Plasm Safeguards Germline Development[J]. eLife Sciences, 2020, 9.

(29) Tatapudy S, Peralta J, Nystul T. Distinct roles of Bendless in regulating FSC niche competition and daughter cell differentiation. Development. 2021 Nov 15;148(22):dev199630. doi: 10.1242/dev.199630. Epub 2021 Nov 22. PMID: 35020878; PMCID: PMC8645206.

(30) Rust K, Byrnes L E, Yu K S, et al. A Single-cell Atlas and Lineage Analysis of the Adult Drosophila Ovary[J]. Nature Communications, 2020, 11(1).

(31) Silver D L, Montell D J. Paracrine Signaling Through the JAK/STAT Pathway Activates Invasive Behavior of Ovarian Epithelial Cells in Drosophila.[J]. Cell, 2002, 107(7):831–841.

(32) Sun, J. Notch-Dependent Downregulation of the Homeodomain Gene cut is Requiredfor the Mitotic Cycle/Endocycle Switch and Cell Differentiation in Drosophila Follicle Cells[J]. Development, 2005, 132(19):4299–308.

(33) Chaddad, Fabio. Drosophila sosie Functions with βH-Spectrin and Actin Organizers in Cell Migration, Epithelial Morphogenesis and Cortical Stability[J]. Biology Open, 2012, 1(10).

(34) Tzolovsky G, Deng W M, Schlitt T, et al. The Function of the Broad-Complex During Drosophila Oogenesis[J]. Genetics, 1999, 153(3):1371–1383.

(35) Jevitt A, Chatterjee D, Xie G, et al. A Single-cell Atlas of Adult Drosophila Ovary Identifies Transcriptional Programs and Somatic Cell Lineage Regulating Oogenesis[J]. PLoS Biology, 2020, 18(4):e3000538.

(36) Li H, Janssens J, De Waegeneer M, et al. Fly Cell Atlas: A Single-nucleus Transcriptomic Atlas of the Adult Fruit Fly. Science. 2022 Mar 4;375(6584):eabk2432. doi: 10.1126/science.abk2432. Epub 2022 Mar 4. PMID: 35239393; PMCID: PMC8944923.

(37) Perinthottathil S, Kim C. Bam and Bgcn in Drosophila Germline Stem Cell Differentiation[J]. Vitamins & Hormones, 2011, 87:399–416.

(38) Clough E, Tedeschi T, Hazelrigg T. Epigenetic Regulation of Oogenesis and Germ Stem Cell Maintenance by the Drosophila Histone Methyltransferase Eggless/dSetDB1[J]. Developmental Biology, 2014, 388(2):181–191.

(39) Xie T, Spradling A C. decapentaplegic Is Essential for the Maintenance and Division of Germline Stem Cells in the Drosophila Ovary[J]. Cell, 1998.

(40) Buszczak M, Paterno S, Spradling A C. Drosophila Stem Cells Share A Common Requirement for the Histone H2B Ubiquitin Protease Scrawny[J]. Science, 2009, 323(5911).

(41) Page S L, Khetani R S, Lake C M, et al. corona is Required for Higher-Order Assembly of Transverse Filaments into Full-Length Synaptonemal Complex in Drosophila Oocytes[J]. Plos Genetics, 2008, 4(9):e1000194.

(42) Witt E, Benjamin S, Svetec N, et al. Testis Single-cell RNA-seq Reveals the Dynamics of de novo Gene Transcription and Germline Mutational Bias in Drosophila[J]. eLife, 2019, 8:-.

(43) Ohlstein B,, Lavoie C A, Vef O,, et al. The Drosophila Cystoblast Differentiation Factor, Benign Gonial Cell Neoplasm, is Related to DExH-box Proteins and Interacts Genetically With bag-of-marbles[J]. Genetics, 2000, 155(4):1809.

(44) Steinhauer W R, Kalfayan L J. A Specific Ovarian Tumor Protein Isoform is Required for Efficient Differentiation of Germ Cells in Drosophila Oogenesis.[J]. Genes & Development, 1992, 6(2):233–43.

(45) Fu Z, Geng C, Wang H, et al. Twin Promotes the Maintenance and Differentiation of Germline Stem Cell Lineage through Modulation of Multiple Pathways[J]. Cell Reports, 2015, 13(7):1366–1379.

(46) Vidaurre V, Chen X. Epigenetic Regulation of Drosophila Germline Stem Cell Maintenance and Differentiation[J]. Developmental Biology, 2021, 473.

(47) Blatt P, Martin E T, Breznak S M, et al. Post-transcriptional Gene Regulation Regulates Germline Stem Cell to Oocyte Transition During Drosophila Oogenesis[J]. Current Topics in Developmental Biology, 2019.

(48) Rastegari E, Kajal K, Tan B S, et al. WD40 protein Wuho Controls Germline Homeostasis via TRIM-NHL Tumor Suppressor Mei-p26 in Drosophila[J]. Development, 2020, 147(2):dev182063.

(49) Casale A M, Cappucci U, Pia Ce Ntini L. Unravelling HP1 Functions: Post-transcriptional Regulation of Stem Cell Fate[J]. Chromosoma, 2021:1–9.

(50) Rhiner C, Diaz B, Portela M, et al. Persistent Competition Among Stem Cells and Their Daughters in the Drosophila Ovary Germline Niche.[J]. Development, 2009, 136(6):995.

(51) Ji Y, Tulin A V. Poly(ADP-ribose) Controls DE-cadherin-dependent Stem Cell Maintenance and Oocyte Localization[J]. Nature Communications, 2012, 3:760.

(52) Yusuke O, Tomoya M, Kosuke A, et al. Defective Autophagy in Vascular Smooth Muscle Cells Enhances Cell Death and Atherosclerosis[J]. Autophagy, 2018:15548627.2018.1501132.

(53) Ponnusamyb M P. Stem Cell Research and Cancer Stem Cells[J]. Journal of Tissue Science & Engineering, 2011, 02(3).

(54) Andersen P R, Tirian L, Vunjak M, et al. A Heterochromatin-dependent Transcription Machinery Drives piRNA Expression[J]. Nature, 2017, 549(7670):54.

(55) Scholz H, Franz M, Heberlein U. The Hangover Gene Defines A Stress Pathway Required for Ethanol Tolerance Development[J]. Nature, 2005, 436(7052):845–7.

(56) Xie T, Spradling A. The Drosophila Ovary: An in vivo Stem Cell System. In Stem Cell Biology (ed. D. R. Marshak R. L. Gardner and D. Gottlieb), 2001, pp. 129–148. New York: Cold Spring Harbor Laboratory Press.

(57) Pearson J R, Federico Z, Laura T G, et al. ECM-Regulator timp is Required for Stem Cell Niche Organization and Cyst Production in the Drosophila Ovary[J]. PLOS Genetics, 2016, 12(1):e1005763

(58) A Díaz-Torres, Rosales-Nieves A E, Pearson J R, et al. Stem Cell Niche Organization in the Drosophila Ovary Requires the ECM Component Perlecan[J]. Current Biology, 2021(51).

(59) Schwientek T, Mandel U, Roth U, et al. A Serial Lectin Approach to the Mucin-type O-glycoproteome of Drosophila melanogaster S2 Cells[J]. PROTEOMICS, 2007, 7(18).

(60) S Shapiro S D. Matrix Metalloproteinase Degradation of Extracellular Matrix: Biological Consequences[J]. Current Opinion in Cell Biology, 1998, 10(5):602–608.

(61) Stetler-Stevenson, W. G. Tissue Inhibitors of Metalloproteinases in Cell Signaling: Metalloproteinase-Independent Biological Activities[J]. Science Signaling, 2008, 1(27):re6.

(62) J Shi, Jin Z, Yu Y, et al. A Progressive Somatic Cell Niche Regulates Germline Cyst Differentiation in the Drosophila Ovary[J]. Current Biology, 2020, 31(4).

(63) Xie T. Control of Germline Stem Cell Self-renewal and Differentiation in the Drosophila Ovary: Concerted Actions of Niche Signals and Intrinsic Factors[J]. Wiley Interdisciplinary Reviews: Developmental Biology, 2013, 2(2).

(64) Trupti P, Chen X, Ekaterina A, et al. Specification and Spatial Arrangement of Cells in the Germline Stem Cell Niche of the Drosophila Ovary Depend on the Maf Transcription Factor Traffic jam[J]. Plos Genetics, 2017, 13(5):e1006790.

(65) Slaidina M, Gupta S, Lehmann R. A Single Cell Atlas Reveals Unanticipated Cell Type Complexity in Drosophila Ovaries [J]. Genome research, 2021, 31(10):1938–1951.

(66) Hinnant T D, Alvarez A A, Ables E T. Temporal Remodeling of the Cell Cycle Accompanies Differentiation in the Drosophila Germline[J]. Developmental Biology, 2017:S0012160617300763.

(67) Kahney E W, Snedeker J C, Chen X. Regulation of Drosophila Germline Stem Cells[J]. Current Opinion in Cell Biology, 2019, 60:27–35.

(68) Perez-Martinez L, Jaworski DM. Tissue Inhibitor of Metalloproteinase-2 Promotes Neuronal Differentiation by Acting as An Anti-mitogenic Signal. Journal of Neuroscience 2005;25:4917–4929.

(69) Ahonen M, Poukkula M, Baker AH, Kashiwagi M, Nagase H, Eriksson JE, Kahari VM. Tissue Inhibitor of Metalloproteinases-3 Induces Apoptosis in Melanoma Cells by Stabilization of Death Receptors. Oncogene 2003;22:2121–2134.

(70) Nemeth JA, Rafe A, Steiner M, Goolsby CL. TIMP-2 Growth-stimulatory Activity: A Concentration- and Cell Type-Specific Response in the Presence of Insulin. Experimental Cell Research 1996;224:110–115.

(71) Mohammed FF, Pennington CJ, Kassiri Z, Rubin JS, Soloway PD, Ruther U, Edwards DR, Khokha R. Metalloproteinase Inhibitor TIMP-1 Affects Hepatocyte Cell Cycle via HGF Activation in Murine Liver Regeneration. Hepatology 2005;41:857–867.

(72) Wang X, Page-McCaw A. A Matrix Metalloproteinase Mediates Long-distance Attenuation of Stem Cell Proliferation. J Cell Biol. 2014; 206(7):923–36.

